# Milanković cycles and Cultural Evolution as important Catalysts for Hominin Genetic Diversification

**DOI:** 10.64898/2026.09.03.749301

**Authors:** Shih-Wei Fang, Pasquale Raia, Aneesh Sundaresan, Chiara Barbieri, Jiaoyang Ruan, Ali R. Vahdati, Elke Zeller, Christoph Zollikofer, Axel Timmermann

**Author notes:** These authors contributed equally (Pasquale Raia, Aneesh Sundaresan). These authors contributed equally (Chiara Barbieri, Jiaoyang Ruan, Ali R. Vahdati, Elke Zeller, Christoph Zollikofer). Senior author (Christoph Zollikofer, Axel Timmermann).

## Abstract

Climate variability is widely considered a key driver of human evolution, yet the mechanisms linking long-term climate change driven by Milanković cycles to hominin demography and genetic diversity remain elusive. Here we couple an agent-based demographic model, incorporating an idealized genetic marker, cultural dynamics, and a realistic Pleistocene climate framework, to quantify how astronomically forced climate and vegetation variability shaped the genetic structure of African hominins over the past 1.9 Myr. We show that warm early Pleistocene climates sustained a continent-wide network of genetically diverse demes until ∼900 ka. The subsequent decline in atmospheric CO₂, associated cooling, and reduced vegetation during the Mid-Pleistocene Transition triggered widespread demographic collapse, with populations surviving only in southern and eastern Africa. This reorganization in population structure leads to a decline in simulated genetic diversity, consistent in timing with genomic and archeological evidence. Introducing cultural innovations in the model, represented as enhanced carrying capacity, enables hominin populations to recover from this diversity bottleneck, adapt to increasingly harsh late Pleistocene environments and long glacial periods, rapidly expand across Africa during interglacials, and ultimately disperse into Eurasia. Our results identify Milanković-scale climate forcing and cultural growth as important catalysts of hominin population structure and genetic diversification.

**SUMMARY:**

- A new computer model links astronomically-driven climate change, population movement, and heredity to explain key aspects of early human evolution in Africa.
- The model suggests that warm early Pleistocene climates helped maintain a continent-wide, genetically diverse network of hominin populations.
- Around 900,000 years ago, cooling during the Mid-Pleistocene Transition caused a major population collapse and a sharp loss of genetic diversity, leaving survival mainly in eastern and southern Africa.
- Later cultural innovations increased carrying capacity, helping hominins recover, spread across Africa during favorable periods, and eventually disperse into Eurasia.

## INTRODUCTION

Our genus *Homo* mainly evolved during the Pleistocene epoch (2.58–0.0117 million years ago), which saw gradual global cooling and strengthening of glacial–interglacial cycles (Fig. 1a). These long-term climate changes were driven by astronomically forced variations in Earth’s axial precession, tilt, and orbital eccentricity, with periods of ∼21, 41, and ∼100 kyr, respectively^1^, known as Milanković cycles. In response to this forcing, climate conditions varied across Africa (Fig. 1), reshaping food availability and habitat suitability, causing populations to expand in some areas, and contract in others^2–4^. These complex demographic dynamics cannot be directly reconstructed from fossils. However, they may have left signatures in the genomes of living humans and in ancient DNA. Recent genetic studies reveal a complex pattern of repeated gene flow among spatially separated demes^5–9^, as well as the complete loss of diversity in other lineages^10–12^. Conversely, geographic isolation^13,14^ and regional fixation of mutations likely promoted the emergence of distinctive *Homo sapiens* lineages, as well as extinction and speciation events in earlier *Homo* species^15–18^.

**Figure 1.**
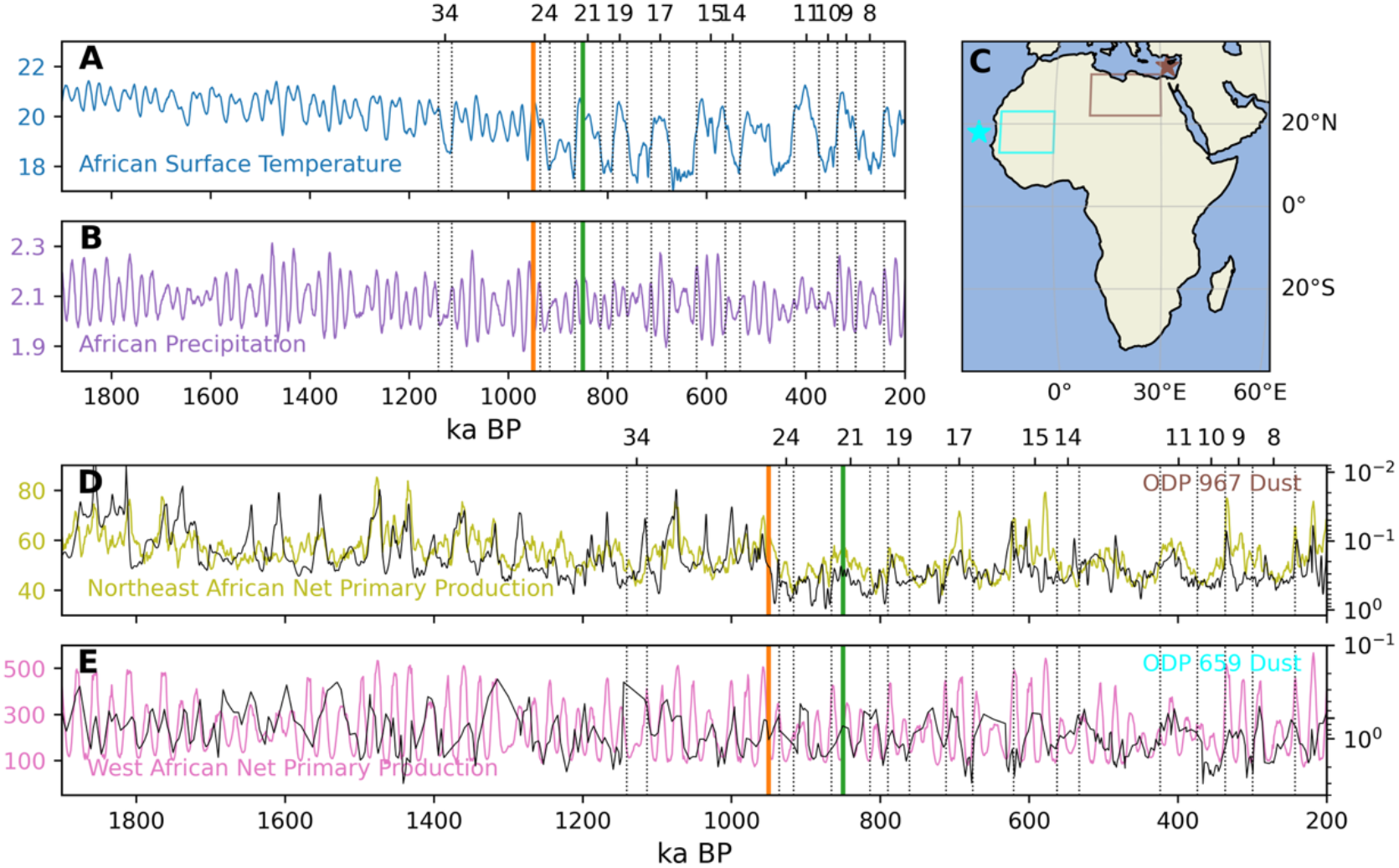
Reconstructed and simulated Milanković-scale climate variability in Africa. **A.** Variations of African Surface temperature [°C] over 40°S-40°N and 15°W-50°E from 3 Ma transient simulation conducted with CESM1.2^29^. **B.** same, but for African precipitation [mm/day]. **D.** Northeast African (26°N-32°N and 20°E-30°E) net primary production [gC/m^2^] with regions close to the dust proxies of Ocean Drilling Program (ODP) site 659^7^. **E.** West African (5°N-15°N and 40°E-50°E) and ODP-967^8^. Dotted lines indicate the important Marine Isotope Stages (MIS) for this study. Orange and green bars highlight the duration of a strong drop in atmospheric CO_2_. **C.** The regions considered and the proxy locations of **D** and **E,** with corresponding proxy colors.

Despite this broad framework being supported by independent evidence, it remains unclear how Milanković forcing and Pleistocene cooling shaped hominin dispersal, interactions, and diversification in Africa. The geographic patterns of lineage diversification and admixture also remain poorly understood because ancient DNA from Africa is still too scarce^19^ and fragmentary to provide a complete picture of our prehistory^19,2^. One way to address this limited phylogeographic resolution, while also testing the impact of Milanković variability on human history, is to use explicit models of human distribution and dispersal. In particular, modern agent-based models^1,20^ can simulate the spread and contraction of multiple populations, their responses to hydroclimate changes, potential admixture events, and the effects of cultural innovation^21–24^ on these processes.

Here, we address these fundamental issues by using the ICCP Climate Agent Hominin Model (ICHAM) (Fig. 2), which combines a spatially explicit paleoclimate-forced agent-based model (ABM)^20,25^, a mitochondrial DNA (mtDNA) inheritance model with a constant mutation rate^26^, and a representation of cultural carrying capacity^27^. We directly examine the continuous population changes in African hominins (including *H. habilis, H. ergaster*, and African *H. heidelbergensis*) and the corresponding shifts in their genetic diversity in response to Pleistocene climate shifts over the past 1.9 Ma. ICHAM simulates virtual individuals^28^ within distinct populations who perform basic activities at an annual time step in a changing environment.

**Figure 2.**
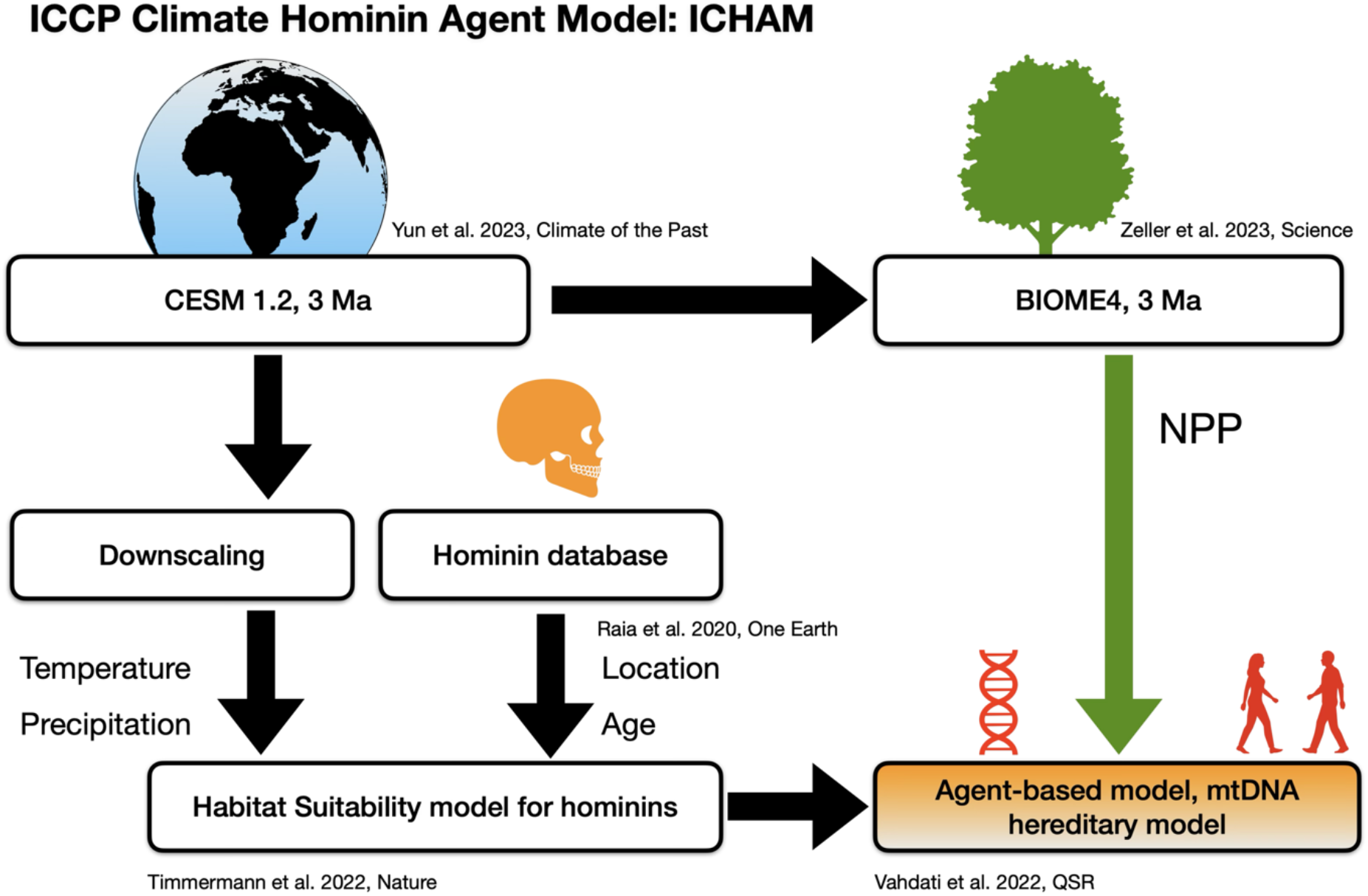
ICCP Climate Hominin Agent Model. Schematics of the ICHAM modeling framework used in this study.

In this study, we chose mtDNA to track ancestry and simulated large-scale population structure over the past 1.9 Ma. Since our model experiments mostly predate the deepest extant observed mtDNA history^30^, the simulated mtDNA statistics cannot be directly compared with empirical data. Instead, the mtDNA from our experiments should be regarded as a means of characterizing how changes in population structure, diversity, and replacement could potentially be stored as genetic information. We acknowledge the limitations of mtDNA as a single genetic locus that is sensitive to drift and incomplete lineage sorting. Its transmission is carried by fewer ancestors than by those who contribute to autosomal genomic inheritance. Nevertheless, due to its non-recombinant nature, small genomic size, and relatively stable mutation rate, mtDNA is particularly suitable for demographic simulations when a controlled set of parameters or scenarios are tested^31^. Moreover, implementing a full forward autosomal genomic inheritance model for the ABM would exceed the available computational capabilities.

## MATERIALS AND METHODS

### ICCP Climate Agent Hominin Model (ICHAM)

Here, we introduce the key components of the ICHAM (Fig. 1). ICHAM consists of an agent-based model^20^ that simulates the life cycle of individual male and female hominins and their maternally inherited mtDNA sequences. The agents are born in one place, grow up, reach sexual maturity, consume food, reproduce, move across the landscape, and die. Hominin food resources are calculated from a carrying capacity, which is a product of two key components: i) land surface NPP obtained from a 3 Ma BIOME4 simulation^24^, and ii) time-evolving continuous habitat suitability estimates^20,25^ for African hominins^15,18^ (Figs. 3a and S1) from 1.9 Ma to 0.2 Ma calculated from a human presence dataset^15,22^ and a realistic transient previously validated paleo-climate model simulation which covers the past 3 Ma^29^. From the geographic positions and mtDNA of individual human agents, we calculate their ensemble mean population density (Fig. 3c) and structure, as well as their metapopulation genetic diversity (Fig. 3b) ^32^.

**Figure 3.**
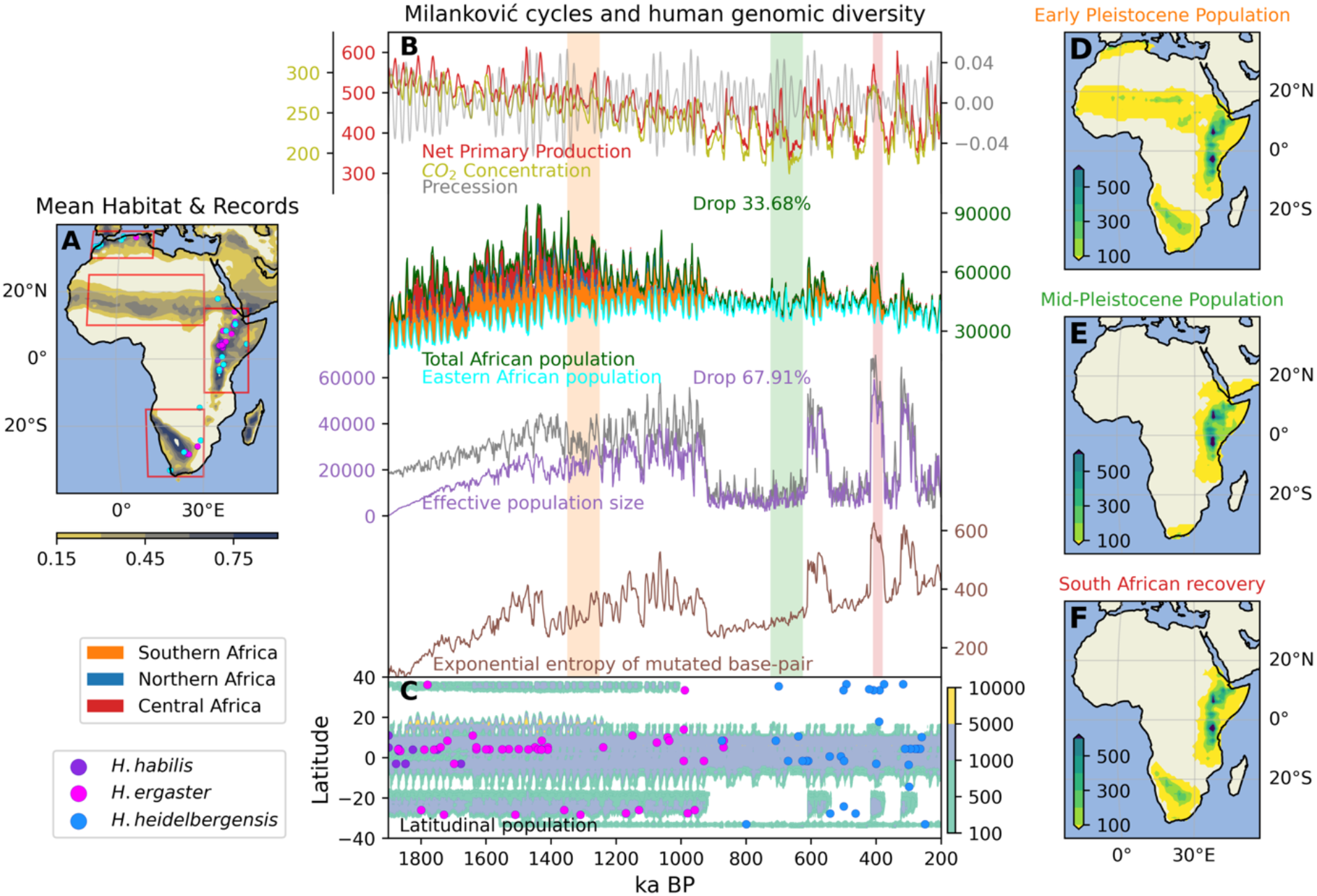
Climate impacts on genetic diversity during the Pleistocene. **A.** Mean habitat suitability of African hominins over the simulation period (1.9 to 0.2 Ma). The dots represent the locations of fossil/archeological records^15^ used in this study: blue-violet for *Homo habilis*, magenta for *Homo ergaster*, and cyan for *Homo heidelbergensis*. **B**. Net primary production [gC/m^2^] (red) variation in Africa (over 40°S-35°N and 15°W-50°E), CO_2_ concentrations [parts per million] (olive), and precession parameter changes (gray) of the 3 Ma simulation^29^. Total African population (dark green), eastern Africa population (cyan), effective population size (red; Methods); and exponential entropy of mtDNA nucleotide diversity^37^ (brown; Methods) are shown. The dark gray line represents the effective population size of a simulation with a longer genetic burn-in time of 300 kyrs. The bars between total African population and eastern African population represent the populations in southern (orange), northern (blue), and central (red) Africa. The percentages are calculated by the ratio between the mean after (green bar from 0.85-0.75 Ma) and before (orange bar from 1.05-0.95 Ma) the 0.9 Ma bottleneck. **C.** Latitudinally aggregated populations of the ICHAM simulation and latitude-time distribution of fossil/archeological records^15^ **D**. Map of averaged population density during the Early Pleistocene (orange bar in B), **E.** after the MPT (green bar in B), and **F.** during the interglacial period MIS 11(red bar in B).

### The Agent Based Model

The ABM in this study is a new implementation of a previously published ABM^20,25^ written in Julia. We track the agent’s mtDNA, consisting of a nucleotide sequence of 16,569 base pairs (bp). For the mtDNA initialization of the ABM we simulate 4,000 individuals in three geographically separated subpopulations: northern, eastern and southern Africa. All individuals start with the same mtDNA sequence and with equal initial probability of the nucleotides A, G, C, and T. We conducted another sensitivity experiment with a 300-kyr-longer burn-in time, allowing a higher starting population size (dark gray line in Fig. 3b) to investigate the impact of genetic equilibration.

The climate forcing for the ABM and habitat model (Fig. 3b), which enters the calculation of the time-varying carrying capacity of the ABM, is based on a 3-million-year (3 Ma) transient climate model simulation conducted with the CESM1.2 model^29^. The 3 Ma simulation uses estimates of greenhouse gas concentrations (CO_2_, N_2_O, and CH_4_)^33,34^, model-based estimates of Northern Hemisphere ice-sheet topography and albedo^33^, and astronomical (Milanković) forcing representing the time-latitude variations of incoming solar radiation^35^. The horizontal resolution of the CESM1.2 simulation is ∼3.75×3.75 degrees in the atmosphere and ∼3 degrees in the ocean. The 1000-year-averaged surface temperature and total precipitation data were downscaled to a 1×1-degree grid and combined with hominin presence data^15,22^ to calculate habitat suitability. NPP is obtained from the well-validated 3 Ma BIOME4 model^24^, which was forced with the transient climate model data of the 3 Ma CESM1.2.

The ABM is run with a yearly timestep, and climate conditions update the carrying capacity, habitat suitability, and biome preference probabilities every 1,000 years, following the overall trajectory of the 3 Ma CESM1.2 climate model simulation^29^. The model domain covers Eurasia and Africa with a horizontal resolution of 25 km. The fixed topography is based on the 5’ Gridded Global Relief Data (ETOPO5)^36^, and the boundary of the continent and water regions, including ocean, lakes, and rivers (areas of increased carrying capacity), are obtained from the Nature Earth Database (https://www.naturalearthdata.com/). The detailed parameters for modeling African hominins, such as birth rate, fertility age, and maximum age, are listed in Table S1.

### Hominin habitat suitability model

The statistical habitat suitability model used in this study is a modified version of the statistical Mahalanobis distance-based climate envelope model (MD-CEM)^18^. Rather than stratifying the MD-CEM for different human species (*H. habilis*, *H. ergaster*, and *H. heidelbergensis* in Africa) by calculating the climate envelope for all fossil records at once^18^, we treat the three African hominins as a time-evolving phylogenetic continuum (Fig. S1) and compute climate envelopes within a rolling window, which includes 40 hominin presence data points. The flexible time range for each window is determined by the estimated minimum/maximum ages across the 40 hominin presence records (30 and 50 records per window show similar variability). Given, that each chunk only includes 40 samples, we restrict the controlling environmental parameters to only a 4-dimensional manifold based on annual mean temperature, annual mean precipitation, annual minimum precipitation (from the 3 Ma CESM1.2 simulation), and terrestrial NPP (from BIOME4) with 1,000-year downscaled averages (1° × 1° horizontal resolution). The habitat suitability model captures crucial aspects of hominin ecology, including food availability and key biophysical factors such as temperature and hydroclimate tolerances.

### Genetic diversity analysis and phylogenetic tree reconstruction

The most relevant parameter extracted from the mtDNA variation of individual agents simulated by the ABM is the effective population size (*N_e_*), which quantifies the number of individuals in a population that contribute genes to the next generation. *N_e_* is estimated after the simulations from the nucleotide diversity ( *π*) with the formula *π* = 2*N_fe_μ*, where *μ* is the mutation rate per generation used in the ABM simulation. The nucleotide diversity is calculated as the mean of the number of site differences in pairwise comparisons 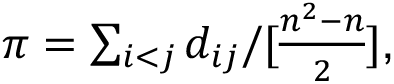 where *di*j is the number of different sites between sequence *i* and *j*, and *n* is the number of females considered, which we chose to be 100 for every 1,000 simulation chunk. Using this subsampling of the total population, we then simply assume the effective population size as twice the size of diverse females, *N_e_* = 2*N_fe_*. We also use two other methods to estimate the effective population size. One is based on the Bayesian Skyline computations of the BEAST model, which derives *N_e_* from the variations in coalescent rates (Fig. 4B), and which we also use to estimate the structure of the phylogenetic tree^38^. Another method is the Watterson estimator^39^, *θ*, which theoretically gives the same number as the nucleotide diversity, i.e., *θ* = 2*N_fe_μ*. The Watterson estimator is calculated from the expected number of segregating sites, *E*(*S*) = *a*_1_*θ*, where 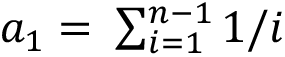 and *S* denote the number of segregating sites, and E refers to the statistical expectation value. Although the values differ, the simulated changes in effective population size are qualitatively very similar across all methods (Fig. 4). The large temporal variations in effective population size derived from the Watterson estimator depend on the number of the small southern African populations within the 100 random samples selected in the calculations. After checking the robustness of the different methods in producing similar outcomes, in particular with regard to the changes during the Mid-Pleistocene Transition (MPT) 1,000-900 ka (Fig. 4), and comparing the results with the statistics of the raw mtDNA data of the ABM, we decided to focus on nucleotide diversity for the main analysis (Fig. 3).

**Figure 4.**
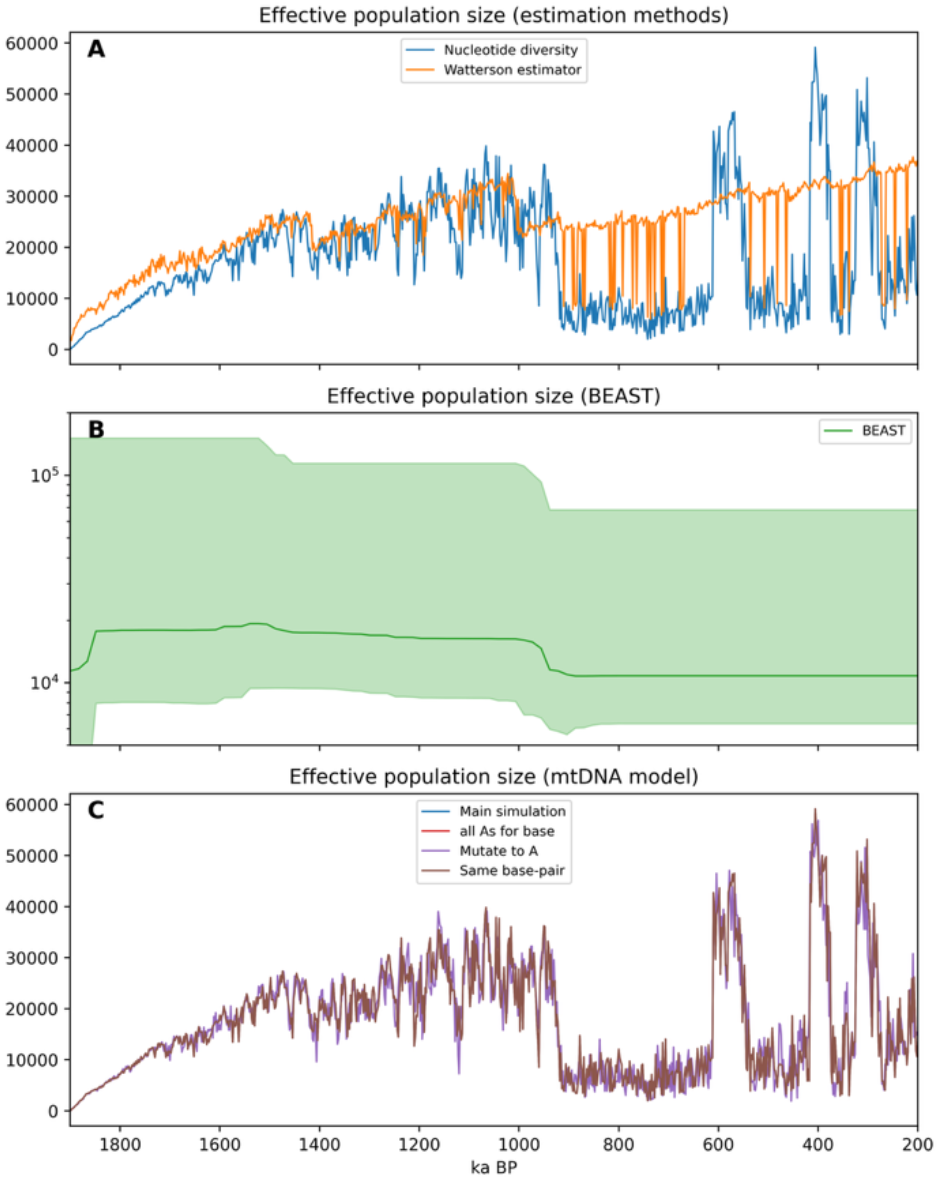
Estimates of effective population size using different methods. **A.** Temporal variations of effective population size calculated by nucleotide diversity (blue) and Watterson estimator (orange). **B.** the BEAST model (green). The green shading is a conservative estimate of the uncertainty calculated from the BEAST model. **C.** Temporal variations of effective population size from random base mtDNA with A-G and C-T mutation (blue; main simulation), from the all As base mtDNA with A-G mutation (red), from random base mtDNA with all-to-A mutation (purple), and from the simulation allowing mutation at the same site (brown).

Multiple methods are used to assess the robustness and underlying causes of the main decline in genetic diversity during the MPT. From a nucleotide composition perspective, we tested whether the main results are affected by the distribution of nucleotide types by setting the initial mtDNA base to all A’s. From the mutation perspective, we tested whether allowing multiple mutations to occur at the same base pair made a difference. We also tested the case that all mutations can only mutate to the same label, A. All these sensitivity experiments reveal very similar changes in genetic diversity (Fig. 4C). We have also subsampled different numbers of female individuals, with virtually no impact on the overall results (not shown).

We also applied multiple methods to estimate the genetic distance between pairs of individuals, the relationships between genetic and geographic distance, and to calculate the genetic contributions of subpopulations, and the diversity of mutated base-pairs (Fig. 5B and C). To this end, we calculated pairwise genetic distance by the Tamura-Nei distance^37^ with the Matlab function seqpdist (Fig. 5A). The genetic diversity represents the mean pairwise distance, which shows a pattern similar to that of the effective population size. The relationship between genetic and geographic distance is calculated using the Mantel test (Fig. S2), which characterizes the correlation coefficient between Tamura-Nei distance and geographic (Haversine) distance. The fixation rate is used to estimate the contribution of subpopulations to metapopulation diversity (Fig. 5D), and it is calculated as (*d_M_* − (∑ *d_s_*/*n_s_*))/*d_M_*, where *d_M_* is the genetic distance across the metapopulation, *d_s_* is the genetic distance within subpopulation *s*, and *n_s_* is the number of subpopulations. To further investigate how base-pair differences impact metapopulation diversity, we calculate the exponential entropy of mutated nucleotides (i.e., the probability of mutated individuals for base-pair) (Fig. 3B). High values of exponential entropy indicate more evenly distributed base-pairs, which includes two perspectives: 1) more random accumulation of mutations to increase the evenness, and 2) more different mutated base-pairs are found over comparable numbers of subpopulations.

**Figure 5.**
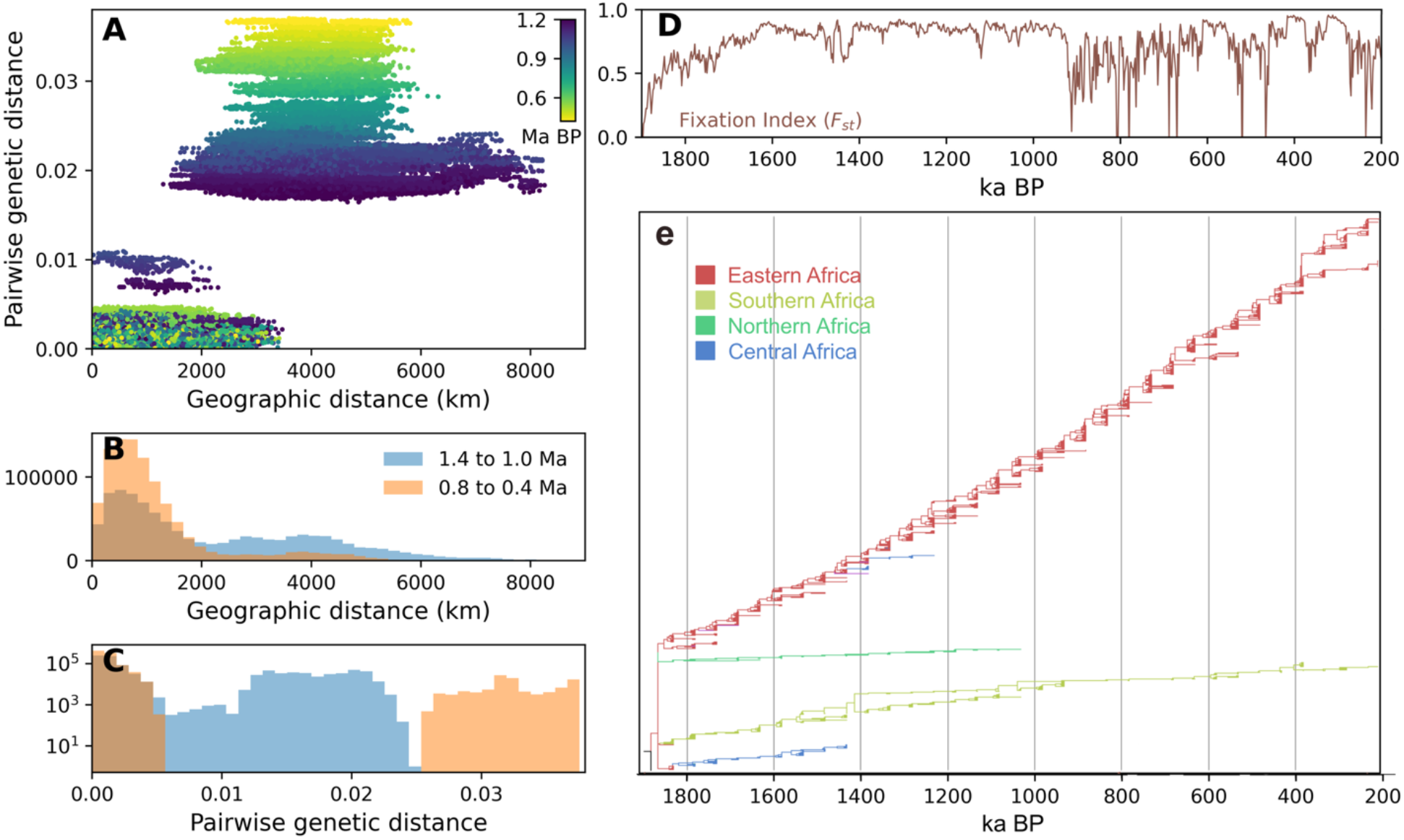
Linking Pleistocene genomic diversification to population structure. **A.** Relation between geographic distance (km) and genetic distance using pairs over 100 randomly selected individuals every 20,000 years from 1.2 to 0.4 Ma. The color indicates age. **B.** Histogram of geographic distance for pairs from 100 individuals each 2,000 years aggregated over two periods: 1.4-1.0 Ma and 0.8-0.4 Ma. **C.** Histogram of genetic distances with logarithmic abscissa. **D.** Fixation rate (Fst) calculated with 1,000 samples from geographically separated subpopulations. **E.** Phylogenetic tree estimated with BEAST2 software using the simulated mtDNA of randomly selected 100 individuals every 50,000 years^40^. Colors indicate the geographic origin of the sampled individuals.

For the phylogenetic analysis of the simulated mtDNA sequences (Fig. 5e) we used the Bayesian Evolutionary Analysis by Sampling Trees (BEAST2) software, version 2.6.7 ^38^. We consider the mtDNA of 100 randomly sampled females in our simulation every 50 thousand years from 1.9 Ma to 0.2 Ma using their age as a tip-date. Only one sample with the starting mtDNA sequence is provided for the root of the tree at 1.9 Ma (corresponding to the most common ancestor). We used a strict clock with a clock rate of about 2.74×10^-8^ base pairs per year^26^, similar to the settings of the simulations. We chose a JC69 model ^41^ and a coalescent Bayesian Skyline^42^ as the tree priors for the simulation. The BEAST calculation is conducted for 15 million generations, and the subsampling is done for every 5,000 generations. The estimated tree file obtained as BEAST output is summarized after discarding the first 20% of the trees (burn-in). We then constructed the Maximum Clade Credibility (MCC) tree using the TreeAnnotator program. The FigTree v1.4.4 software is then used for visualization of the phylogenetic tree (Fig. 2E). The Bayesian skyline plot data (Fig. 5e) is generated using Tracer v1.7.2^43^ and by loading the BEAST output files (log and trees file).

## RESULTS

### The early Pleistocene (1.9-1 Ma)

The time-evolving demographic/genetic hominin simulations are initialized in the early Pleistocene at 1.9 Ma by placing 4,000 individuals with the same base mtDNA in three regions: eastern, southern, and northern Africa. These regions are selected based on records of mid-Pleistocene human presence (*H. ergaster*) during this time (Fig. 3A). During the first 400 kyrs of the simulation, the African population size increases steadily, reaching up to ∼90,000 individuals around 1.5 Ma (Figs. 3B, S3, and S4). The eastern African population, which accounts for >60% of the total *Homo* individuals alive at that time, expands southward, reaching Lake Malawi while one branch moves westward ∼1.8 Ma into an area where it experiences the strong meridional shifts in Africa’s Intertropical Convergence Zone (ITCZ)^18^ with an orbital period of ∼21 kyrs (Figs. 3B and S3), linked to the precession cycle of incoming solar radiation. The southern African population size fluctuates between 10,000 and 20,000 individuals at any given time (Fig. 3B), reaching the maximum extent in the Kalahari region. The northern African population remains confined to the coastal area, counting no more than 8,000 individuals at any given time. The simulated geographic distribution of populations in ICHAM qualitatively matches the spatiotemporal distribution of archeological records (Figs. 3A and C). The spatial separation between the populations in our simulation increases the total metapopulation genetic diversity (Fig. 3B) during the first 500 thousand years (kyrs) of the simulation (1.9-1.4 Ma), as indicated by the estimates of nucleotide diversity (i.e., effective population size)^32^ over 100 randomly selected individuals sampled every 2,000 years (Fig. S4E). These results are robust with respect to reasonable changes in parameters and initial conditions, e.g., with a longer genetic burn-in time of 300 kyrs and with a different base-mtDNA initialization (dark gray line in Figs. 3B, 4C and Fig. S3E).

The eastern African population reaches a peak of some 40,000-50,000 individuals (Figs. 3 and S4A) around 1.4 Ma, with only small precessional-scale (21 kyr) fluctuations (+/-10%), underscoring its role as a climatically stable source area for African hominins. Initially, simulated populations in northern, southern, and eastern Africa remain disconnected, resulting in separate branches in the phylogenetic tree (Fig. 5E), as reconstructed using BEAST^40^. These regional populations do not exchange individuals with different mtDNA with each other, as indicated by the high population distances and the corresponding Fixation Index (Fig. 5D). In our baseline simulation, the central and northwestern African populations become permanently extinct between Marine Isotope Stage (MIS) 34 (∼1.2 Ma) and MIS22 (∼0.9 Ma), respectively (Figs. 5E and S4). Since no abrupt habitat change is simulated in these regions during this period (Figs. S1C-E), the two simulated local extinctions are associated with sharp drops in NPP related to decreasing temperatures and reduced food resources (Fig. 2 and 3B) and a nonlinear, threshold-like demographic response. These climatically driven early hominin depopulation events in central and northern Africa represent reductions of 25% and 10% in total African population size, respectively (Fig. 3B). Their disappearance is related only to a slight decrease in effective population size and nucleotide diversity (Fig. 4), which accounts for both changes in diversity across distributed populations and the gradual accumulation of mutations within each subpopulation. The 20-25% drop in exponential mtDNA entropy (Fig. 3B), around 1.05 Ma, on the other hand, highlights a substantial decline in genetic diversity corresponding to the regional subpopulation extirpation.

### The Middle Pleistocene Transition (1.0-0.774 Ma)

The MPT ∼0.9 Ma marks a shift from warmer early Pleistocene conditions towards colder climates^44^. It was characterized by an intensification of glacial cycles and a subsequent shift in climate from a 41 thousand-year (kyr) pacing, due to changes in Earth’s axis tilt, to an 80-120 thousand-year (kyr) periodicity (Fig. 3B)^33,45^, related to the combined effects of changes in Earth’s orbit and axis wobble. In Africa, paleoclimate simulations and reconstructions, e.g., from marine sediment core ODP967, provide clear evidence for a substantial temperature drop^29^ during the MPT and a trend toward increasing aridity^46–48^ (Fig. 1). More specifically, MIS22 (∼0.9 Ma)^49^ represents the single most pronounced orbital-scale cooling during the early MPT (Figs. 3B and 6D). Across this event, ICHAM simulates a collapse in genetic diversity, with the total effective population size in Africa decreasing from ∼26,000 to just ∼7,000 individuals (72% reduction) (Figs. 3B, 4, and 6E). In contrast to this massive shift in overall genetic diversity, the continental population size decreases by only ∼20%. This decline can be traced to a contraction of the southern African deme to only 1,000 to 2,000 individuals remaining in the very stable coastal habitat around Nelson Mandela Bay (Figs. 3E and S3C). Our results provide evidence for a population decline and sharp genetic transition occurring through the 0.9 Ma MPT, which is qualitatively consistent with the low density of the archeological artifacts across Africa^50,51^ and a previously-reported - yet controversial^52^ - genetic bottleneck from extant DNA data which has been identified using the FitCoal bioinformatic algorithm^53^.

**Figure 6.**
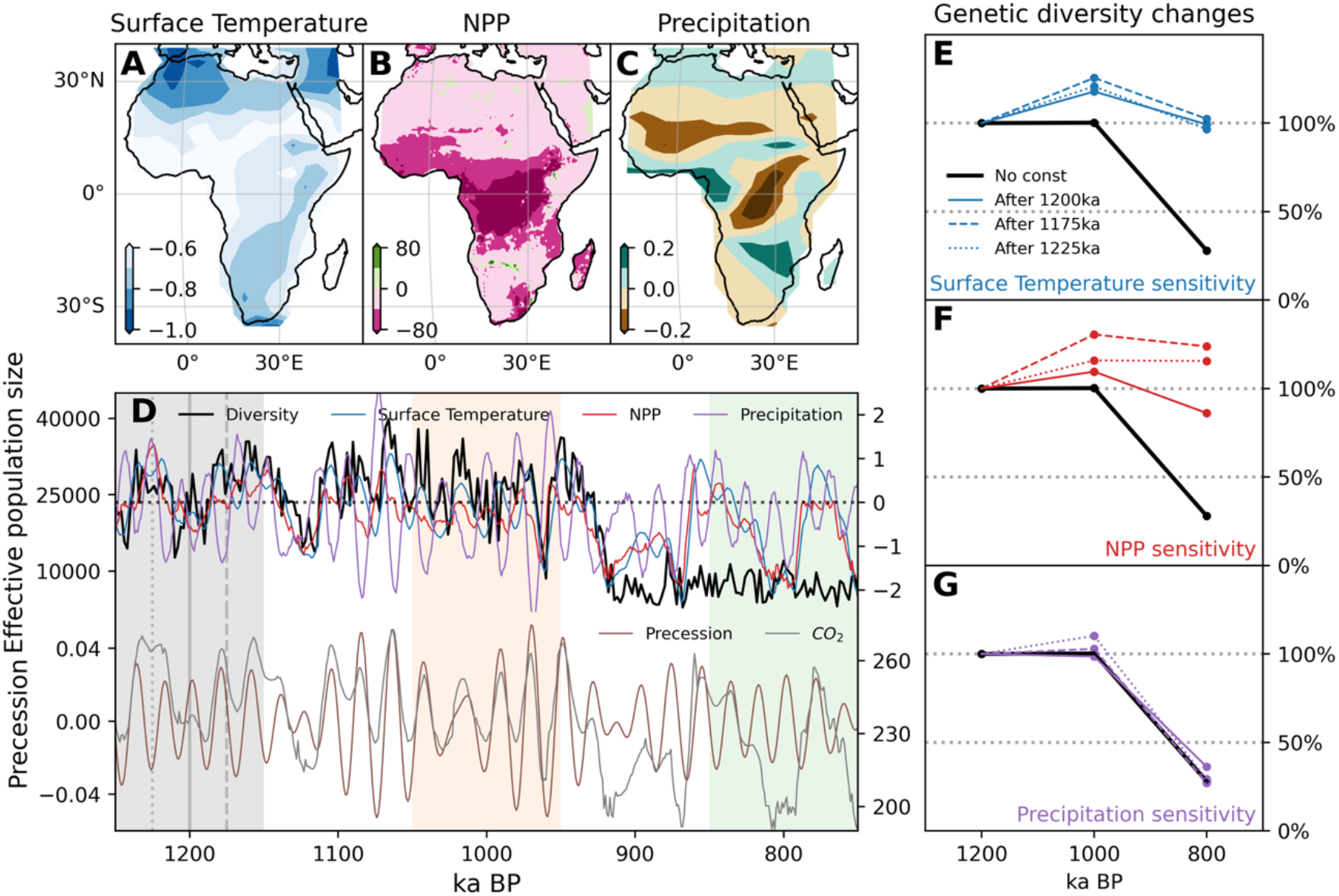
Milanković-scale climate impacts on genetic diversity. **A.** Surface temperature anomaly [°C] across the bottleneck event (0.95-0.85 Ma). Anomalies are calculated with respect to the past 300 kyrs mean (1.25-0.95 Ma). **B.** for net primary production [gC/m^2^/year]. **C.** for precipitation [mm/day]. **D.** Temporal variations of effective population size (diversity; black) of the simulation (upper-left y-axis) and the normalized index of African surface temperature (blue; Fig. 1A), NPP (red; Fig. 2B), and precipitation (purple; Fig. 1B; upper-right y-axis). The lower panel shows precession (brown; left y-axis), and CO_2_ concentration [parts per million] (gray; right y-axis). **E.** Averaged changes in effective population size (genetic diversity) from the constant surface temperature simulations over three periods indicated with shading in d; gray shading for 1.25-1.15 Ma, orange shading for 1.05-0.95 Ma, and green shading for 0.85-0.75 Ma. The percentages are calculated with respect to the 1.2 Ma mean (gray shading). The start time of constant surface temperature for calculating habitat suitability is highlighted with distinct line styles in d (vertical lines) and e, using solid lines for 1.2 Ma, dashed lines for 1.175 Ma, and dotted lines for 1.225 Ma. The original ICHAM simulation is indicated in solid black lines. **F.** for constant NPP simulations, **G.** for constant precipitation simulations.

The fate of individual ICHAM agents includes a random element. To address the effect of stochasticity on the outcome, we repeat the ICHAM simulations 10 times. We note that both central and northern African populations can survive beyond the glacial period MIS34 (1.126 Ma), five and three times, respectively, but never beyond the 0.9 Ma event (Fig. S3). In stark contrast to northern and central Africa, the southern African population goes extinct only once in ten simulations, and in eastern Africa extinction never occurs (Fig. S3). This supports the idea that eastern Africa likely served as a refugium and genetic reservoir for humans, from which subsequent dispersals and evolutionary changes originated, potentially including the transition from *H. ergaster* (i.e. African *H. erectus*) to *H. heidelbergensis*.

In our simulations, the metapopulation genetic diversity depends strongly on the number of individuals and on habitat connectivity among subpopulations, which change in response to the Milanković climate forcing and, more specifically, to the MPT transition. Individual female agents from the same region (within less than ∼2,000 km) (Fig. 5 A, B) maintain approximately a genetic equilibrium with pairwise genetic distances < ∼0.007 throughout the simulation (Fig. 5C). In contrast, the genetic distance between individuals from distinct subpopulations increases over time (Fig. 5A). The simulated genetic collapse at ∼0.9 Ma is related to a contraction of population structure and a corresponding reduction in pairwise geographic distances (Fig. 5B), where most of the pairs originate from the same region - eastern Africa. The strong linkage between genetic and geographic distances throughout most of the transient simulation is further supported by the high Mantel test^54^ statistic (Fig. S2).

To provide deeper insights into the impact of environmental conditions, population size, and genetic diversity, we investigate surface temperature, NPP, and precipitation (i.e., rainfall) across Africa. These variables, which capture the MPT and correlate with the synchronous decline in genetic diversity (Fig, 6D), are driven by combinations of atmospheric CO_2_ forcing and Milanković cycles in shortwave radiation^29^. An Empirical Orthogonal Function (EOF) analysis ^55^ shows that surface temperature and NPP are primarily driven by atmospheric CO_2_ changes (Fig. S5). In contrast, precipitation is more sensitive to the Earth’s axis wobble (precession) (Fig. S6).

The changes in effective population size covary with African surface temperature and NPP (Fig. 6D). Before the MPT, higher genetic diversity typically occurred at higher CO_2_ levels during positive precession (austral summer perihelion). This led to higher temperature and NPP (Fig. 6D) in key areas that contribute the most to the African population growth. During the MPT, Africa experienced a continental-scale cooling which is particularly pronounced in northern and southern Africa (Fig. 6A). Qualitatively consistent with the drying trend observed in paleoclimate reconstructions (Fig. 1D) and the increase in atmospheric dust levels, the model also simulates a drying in sub-Saharan, equatorial and southern Africa through this period (Fig. S5). As a result, NPP, which is a reliable proxy for human food availability (at least outside rainforests) decreased (Fig. 6B), limiting the resources accessible to the virtual agents in ICHAM. Additional sensitivity experiments conducted by holding individual climate parameters constant in the habitat suitability model (Figs. 6E-G) highlight the key role of orbitally controlled changes in temperature and NPP in driving the collapse of genetic diversity.

### The Middle Pleistocene (0.774-0.2 Ma)

Despite considerable changes in NPP during interglacials MIS 21, 19, and 17 (at ∼0.85, ∼0.75, 0.7 Ma, respectively), total and effective population size remain relatively stable after the MPT for at least 300 kyrs (Figs. 3B, 6D), albeit at a lower level than during the early Pleistocene. Only the strong positive NPP and habitat suitability anomalies in South Africa during MIS 15, 11, and 9 (Fig. S1D) (at ∼0.6, 0.4, 0.3 Ma), which create temporary warm and wet corridors into the Kalahari region, lead to population pulses and increased metapopulation genetic diversity (Fig. 3B). According to the simulations, these conditions further allowed small population propagules to survive along the coast of southern Africa (Fig. 3E), to disperse inland, and even to drive the population size back to pre-MPT bottleneck levels. One interesting consequence of these modeling results is that Southern Africa could have potentially been continuously occupied, indicating the presence of stable refugium habitats, with implications for the emergence of later human species, such as later *Homo sapiens*^5,56^.

The evidence presented so far assumes that cultural changes and innovations had no consequences for hominins’ capacity to convert NPP into increases in population size^1^. This is simplistic and at odds with evidence suggesting a major shift in the rate of cultural innovation took place during the Middle Pleistocene^51^ (Fig. 7A). We added an additional layer of realism investigating the potential impact of cultural innovation (Fig. 7A) in ICHAM by adjusting the model parameter (K_max_, see Methods) that controls the maximum number of individuals that can occupy and survive per unit area, thereby assuming that cultural innovation increases carrying capacity globally^57,58^. Note that we do not simulate cultural diffusion here, but rather assume a somewhat idealized spatially independent increase in cultural skills across hominin populations.

**Figure 7.**
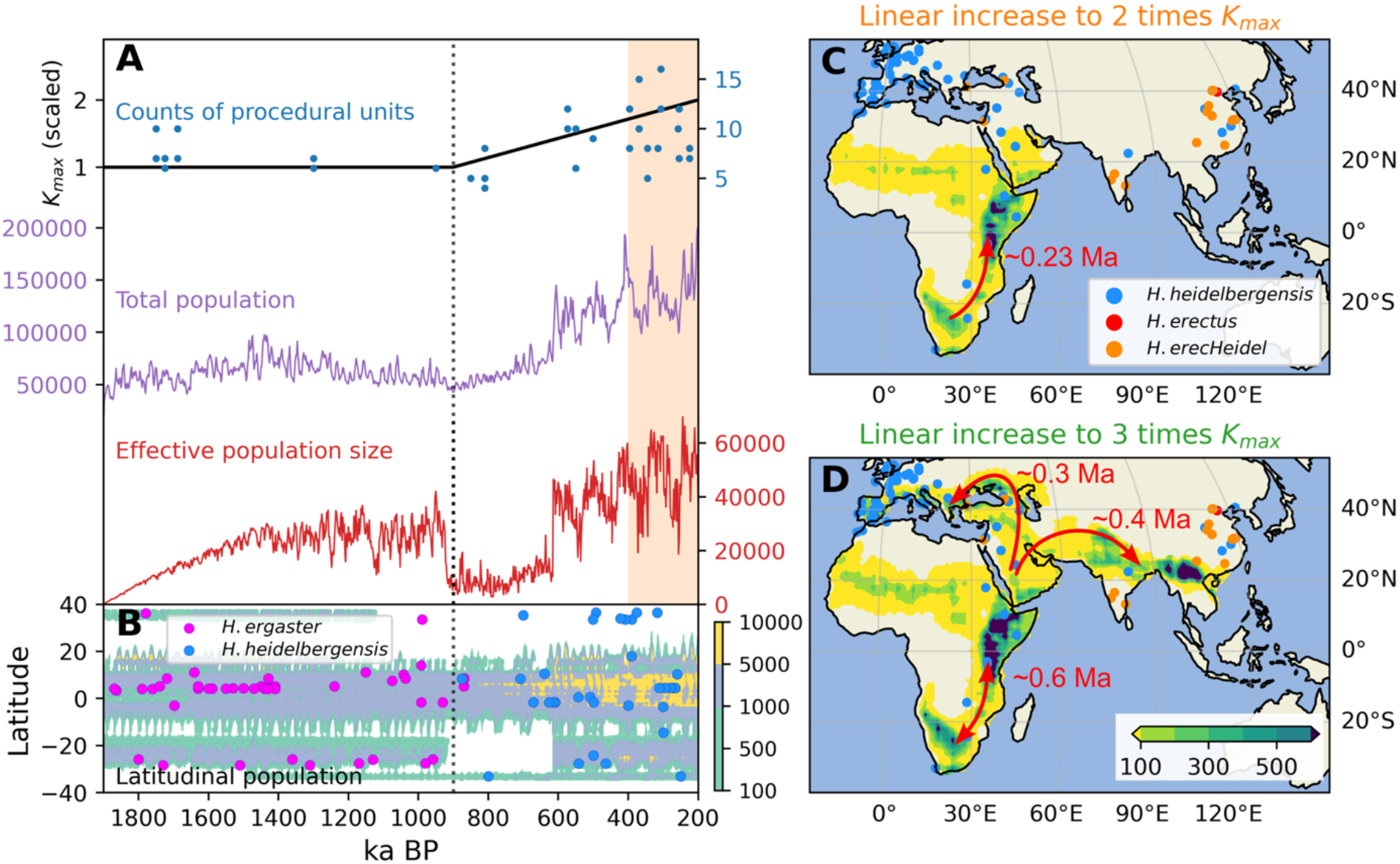
Combined effect of Milanković cycles and cultural evolution on hominin dispersals and population connectivity. **A.** Temporal changes of the K_max_ ABM parameter (black), mimicking cultural development, as indicated by observed counts of procedural units for Africa records (blue dots)^51^. Total population and effective population size of the simulation. **B.** Latitudinally aggregated populations of the ICHAM simulation and latitude-time distribution of fossil/archeological records^15^ **C.** Mean populations over 0.4-0.2 Ma as shown in orange shading in a for the same simulation. The red arrows symbolize the dominant dispersal directions. The timing of the dispersal events is labeled aside. Colored dots indicate the fossil and archeological records of human presence for the period: cyan for *Homo heidelbergensis*, red for *Homo erectus*, and blue for *Homo heidelbergensis/Homo erectus*. **D.** same as B, but for the simulation with a linear increase to tripled K_max_.

We conduct two different types of sensitivity experiments: i) using different yet constant values of K_max_ to examine the idealized impact of cultural adaptation on hominin populations (Figs. S7 and S8), and ii) using linearly increasing K_max_ after the MPT to mimic the observed global trends^51^ in observed tool complexity (Fig. 7A). With constant low levels of cultural adaptation, even at the lowest value of K_max_ tested here, eastern African populations survive the MPT (Fig. S7A), confirming the notion that eastern Africa may have served as the source population for Pleistocene *Homo* in Africa. In contrast to eastern Africa, other regions experience earlier extinctions before the MPT for lower values of K_max_ (Figs. S7B-D), which highlights a stronger human vulnerability to climate conditions in these regions relative to their assumed cultural complexity. Higher K_max_ values also improve survival and connectivity among populations. This situation increases the likelihood of admixture between southern and eastern African populations (Fig. S8 and S9), leading to an evident—yet transient—increase in genetic diversity in the largest eastern African population.

The ICHAM simulations, which assume a moderate two-fold linear increase in cultural complexity post-MPT (Fig. 7A), show a substantial increase in population size, reaching ∼160,000 individuals at the end of the simulation around 0.2 Ma when K_max_ is doubled (Figs. 7A, S9 and S10A-C). The sharp drop in genetic diversity at 0.9 Ma persists for approximately 300 kyrs. During and after the subsequent interglacial event at ∼0.6 Ma, genetic diversity increases continuously, with higher (lower) values for interglacial (glacial) conditions (Fig. 7A). Setting K_max_ to increase up to three times the initial value post-MPT, simulating a substantial cultural impact on population growth, hominins even disperse out of Africa and reach southeastern Europe after 0.3 Ma (Figs. 7D, S9 and 10D-F). These simulated mid-to-late Pleistocene out-of-Africa dispersals are qualitatively consistent with the earliest evidence of *Homo sapiens* in Greece and the Near East^59,60^ (Fig. 7D). The ICHAM simulations further suggest extensive admixture between southern and eastern African populations in the 0.6-0.23 Ma interval (depending on the applied trend in K_max_). The population admixture occurs with individuals from southern Africa dispersing towards eastern Africa (Fig. 7C), which is consistent with the hypothesis that both regions may have contributed to the appearance of *Homo sapiens*^61–63^. Interestingly, recent ancient DNA analysis points to the same biogeographical pattern^56^, albeit for later periods.

## CONCLUSION

We presented explicit modeling evidence that Milanković cycles and the associated changes in food resources, habitats, and subpopulation structure, and an overall growth of cultural carrying capacity, had a fundamental impact on population structure and the genetic diversity of African hominins. The most pronounced signal is observed for the MPT, when the onset of glacial conditions of unprecedented magnitude (MIS22 ∼0.9 Ma) caused a ∼70% loss in nucleotide diversity, as illustrated by independent methods (Fig. 4). A similar, albeit much more pronounced, loss of genetic diversity around 0.93 Ma has been reported based on extant genomic data^53,64^; albeit met with scepticism because of potential drawbacks in the analysis method^52,65,66^. Independent archaeological data (Fig. 7B) further support a period of fewer hominin occurrences across Africa from ∼900-700 ka and a complete absence at latitudes between 30°S and 5°S, which is consistent with the simulated population densities (Fig. 7B).

As the world entered a different climate regime during the MPT, characterized by stronger and longer lasting glacial cycles, surviving hominin populations—generally attributed to the *H. heidelbergensis* lineage—began to show key behavioural and physical shifts including changes in tool production^51^ (Fig. 7A), fire-making^67^, larger brain size^68,69^. These developments may have helped hominins overcome harsher environments, as implied by the arrival of *Homo heidelbergensis* in Britain^22^ ∼600 ka.

Our conclusions rely in part on the transient climate responses of realistic climate^29^ and vegetation models^24^ to a model-based estimate of atmospheric CO_2_ conditions during the MPT until 0.8 Ma^33^ and ice-core data thereafter. Whether atmospheric CO_2_ concentrations dropped sharply during MIS22 remains an open question, but a reconstructed tropical ocean cooling^70^ supports this scenario. More direct evidence is expected from the ongoing Beyond-EPICA ice core initiative, which will soon provide direct measurements of past CO_2_ levels across the entire MPT.

The subject of environmentally-induced human population and genetic bottlenecks has received widespread attention in recent years with cases including i) a depopulation event of Europe around 1.126 Ma due to unprecedented deglacial climate stress^71^ ii) a hypothesized – and heavily debated – close shave of *H. sapiens* triggered by the massive eruption of the Toba super volcano ∼74 kyrs ago^72^, or iii) the recently reported drop in effective population size of all human groups between 0.1-0.02 Ma, with hypothesized potential climatic contributions^73^. Testing these scenarios and quantifying climate causality require multidisciplinary efforts and the integration of independent lines of evidence, including the forward-modeling approach presented here.

## RESOURCE AVAILABILITY

### Lead contact

Further information and requests for resources and reagents should be directed to and will be fulfilled by the lead contact, Shih-Wei Fang.

## ACKNOWLEDGMENTS

This work was supported by the Institute for Basic Science (IBS), Republic of Korea, under IBS-R028-D1. J.R. receives funding from the IBS, Republic of Korea, under IBS-R028-Y3. We acknowledge insightful discussions with Cosimo Posth and Maria Spyrou on the use of the BEAST software.

## AUTHOR CONTRIBUTIONS

Conceptualization, S.W.F. and A.T.; methodology, S.W.F, A.S., C.B., P.R., J.R., A.V., E.Z., C.Z. and A.T.; investigation, S.W.F. and A.T.; writing—original draft, S.W.F. and A.T.; writing—review & editing, S.W.F, A.S., C.B., P.R., J.R., A.R.V., E.Z., C.Z., and A.T.; funding acquisition, A.T.; resources, P.R., J.R., A.R.V., E.Z. and A.T.; supervision, C.B., P.R., C.Z., and A.T.

## SUPPLEMENTARY FIGURES

**Figure S1.**
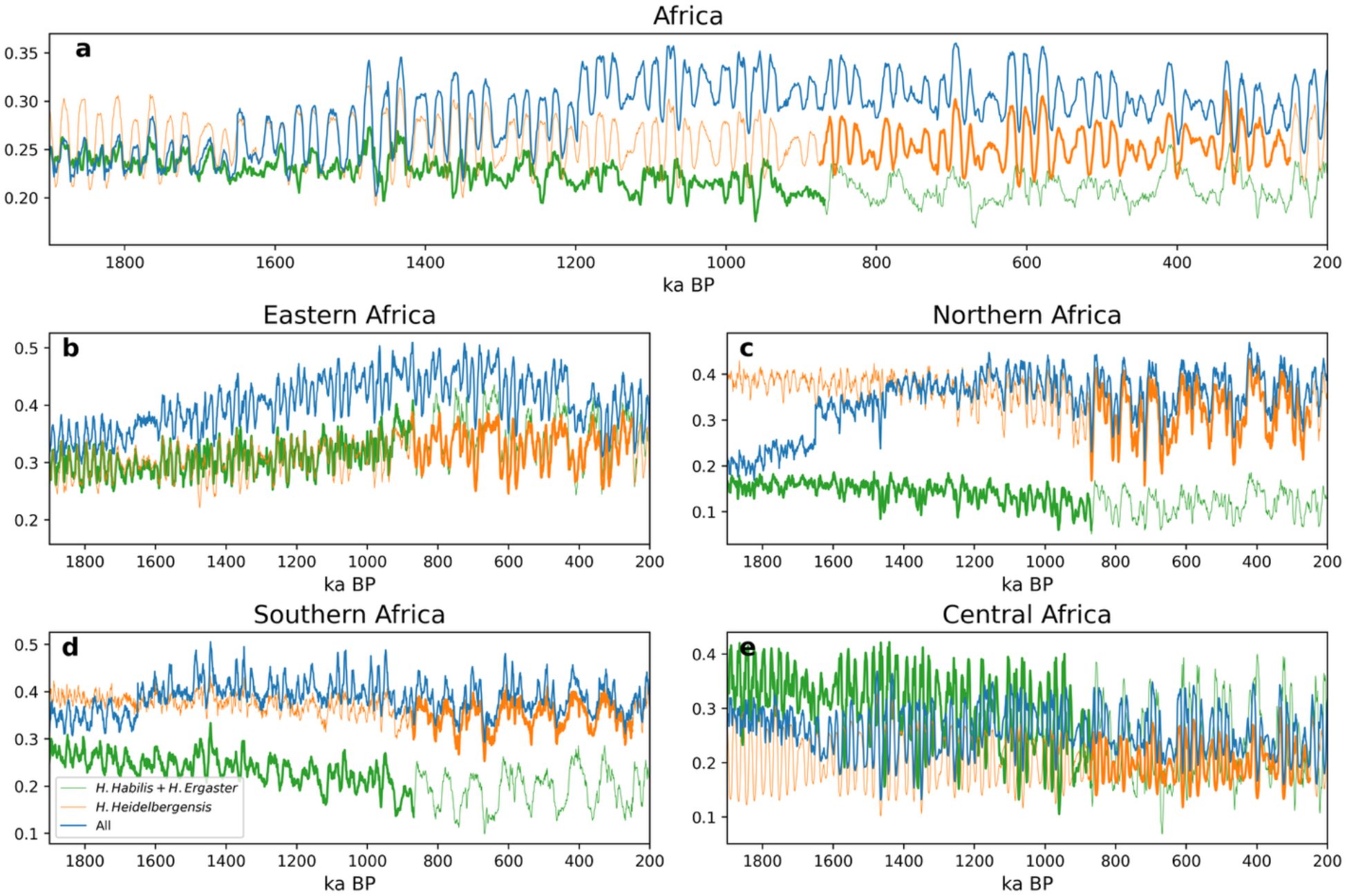
Temporal habitat suitability changes of different considerations of African hominin groups **(a)** Temporal habitat suitability changes in Africa calculated from (green) climate niche models without time-rolling estimated for *Homo habilis* and *Homo ergaster* in Africa, (orange) *Homo heidelbergensis* in Africa, and (blue) climate niche models with time-rolling from all *Homo habilis* and *Homo ergaster* and *Homo heidelbergensis* in Africa. **(b)** for eastern Africa over 10°S-15°N and 30°E-45°E. **(c)** for northern Africa over 30°N-38°N and 10°W-12°E. **(d)** for southern Africa over 35°S-15°S and 10°E-30°E. **(e)** for central Africa over 10°N-25°N and 10°W-30°E.

**Figure S2.**
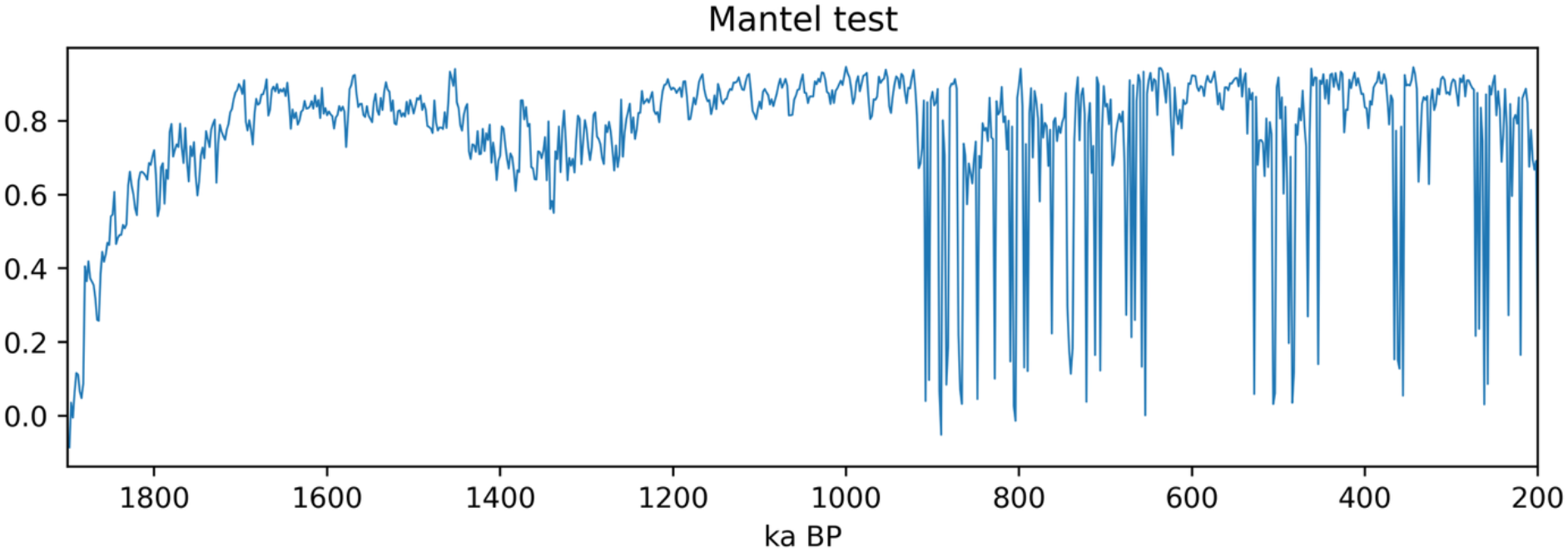
Collapses of high connectivity between genetic and geographical distance after bottleneck. Temporal changes of standardized Mantel test between Nei-Tamura genetic distance and geographic distance of sample pairs.

**Figure S3.**
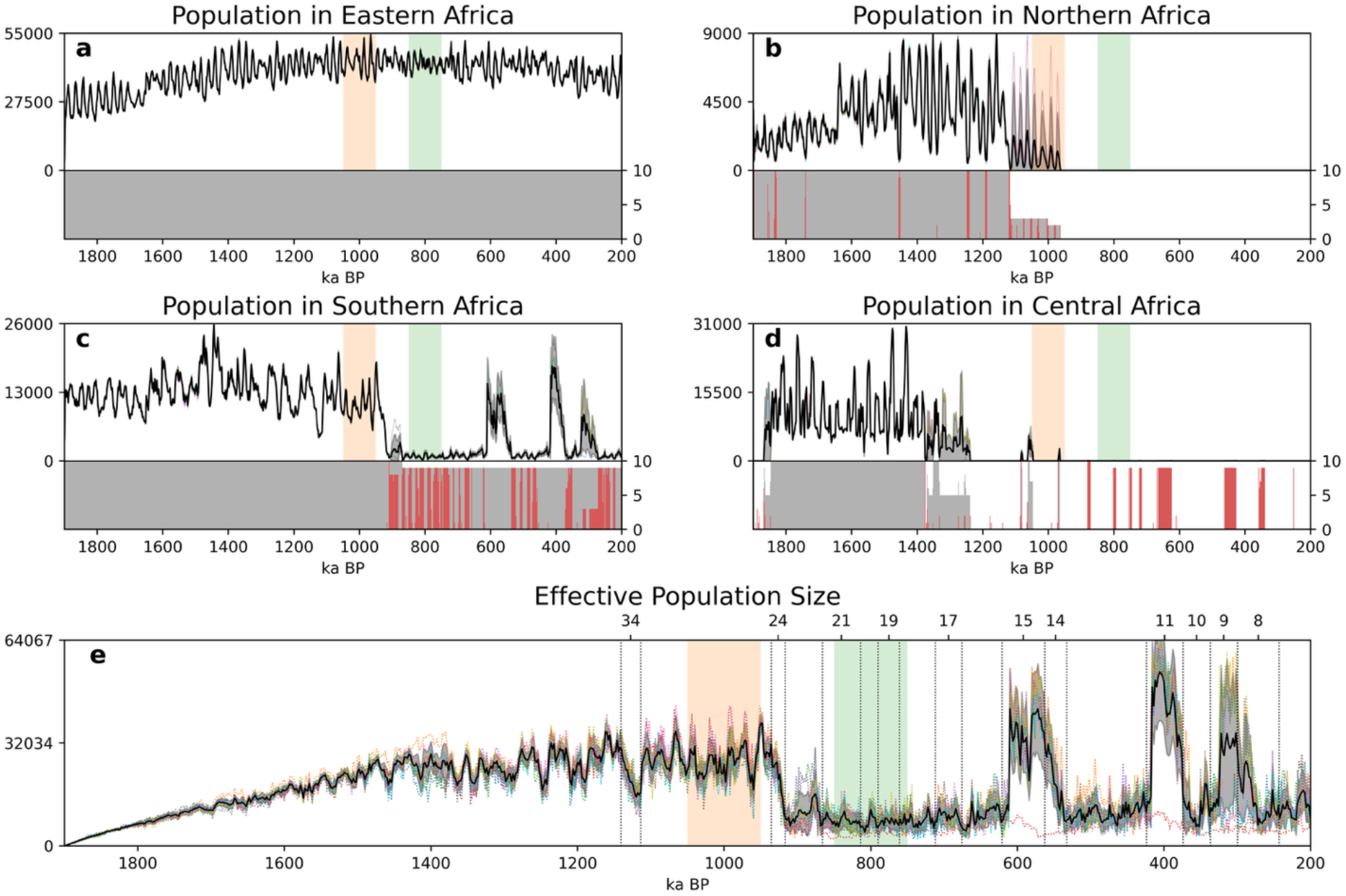
Distinct timing of regional extinctions from 10 ensemble-member simulations. **(a)** Evolution of population (black) in eastern Africa averaged over 10 ensemble members of ICHAM simulations (upper; left y-axis). Individual simulations are highlighted with colored dotted lines, and the spread with one standard deviation is depicted with gray shading. Number of simulations with survivals in eastern Africa over 10 ensemble simulations (lower; right y-axis). **(b)** for northern Africa. The red colors indicate simulations with under 1,000 survivals. **(c)** for southern Africa. **(d)** for central Africa. **(e)** Temporal evolution of effective population size for 10 ensemble members and their mean (thick black). The important Marine Isotope Stages (MIS) for this study are indicated with dotted lines. The orange and green shadings indicate averaging periods before and after 0.9 Ma event calculated in Fig. 1b.

**Figure S4.**
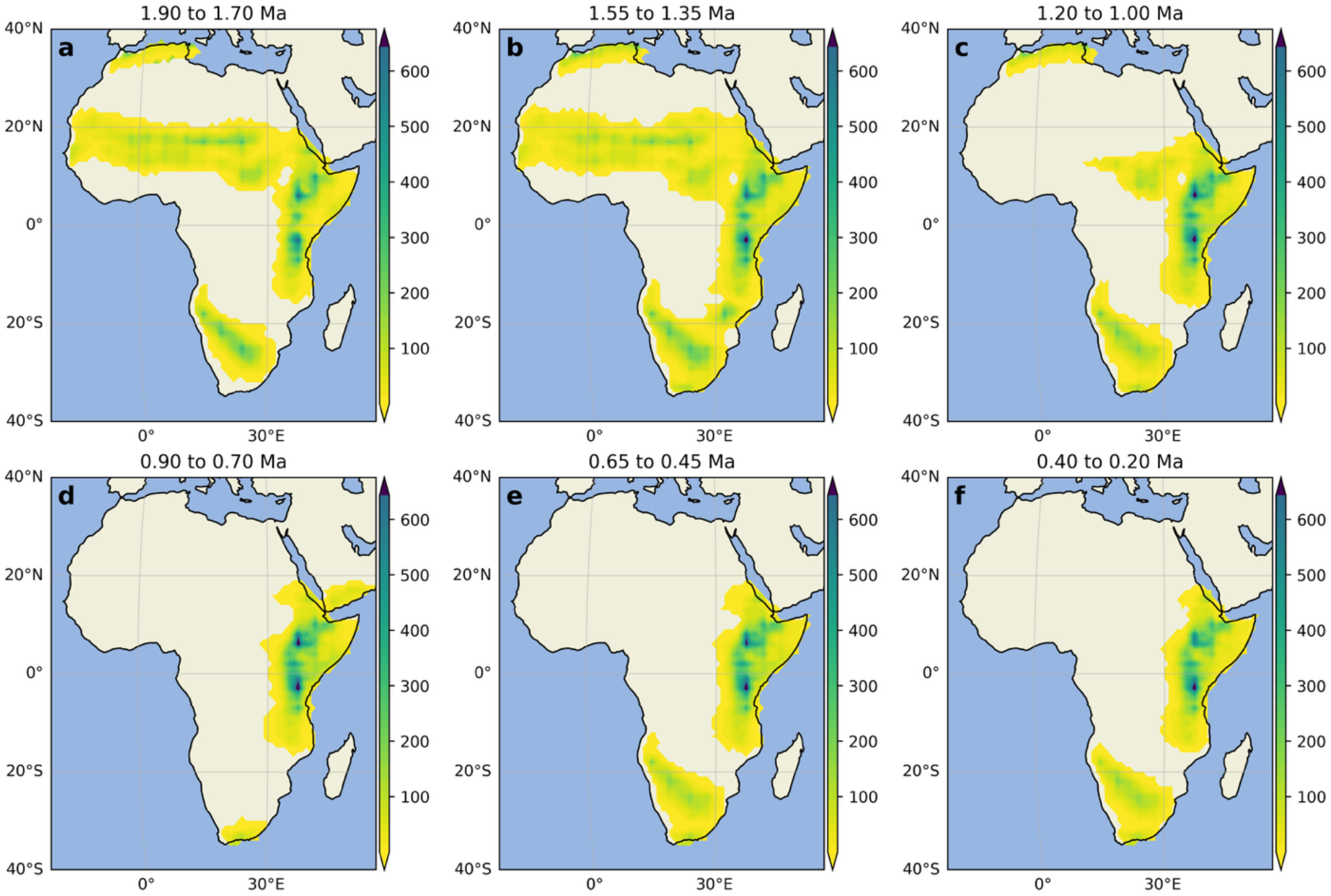
Time slices of population density from the main ICHAM simulation(a) Map of averaged population density (100 km by 100 km) over 1.9 to 1.7 Ma for the main ABM simulation. **(b)** for 1.55 to 1.35 Ma. **(c)** 1.3 to 1.0 Ma. **(d)** 0.9 to 0.7 Ma. **(e)** 0.65 to 0.45 Ma. **(f)** 0.4 to 0.2 Ma.

**Figure S5.**
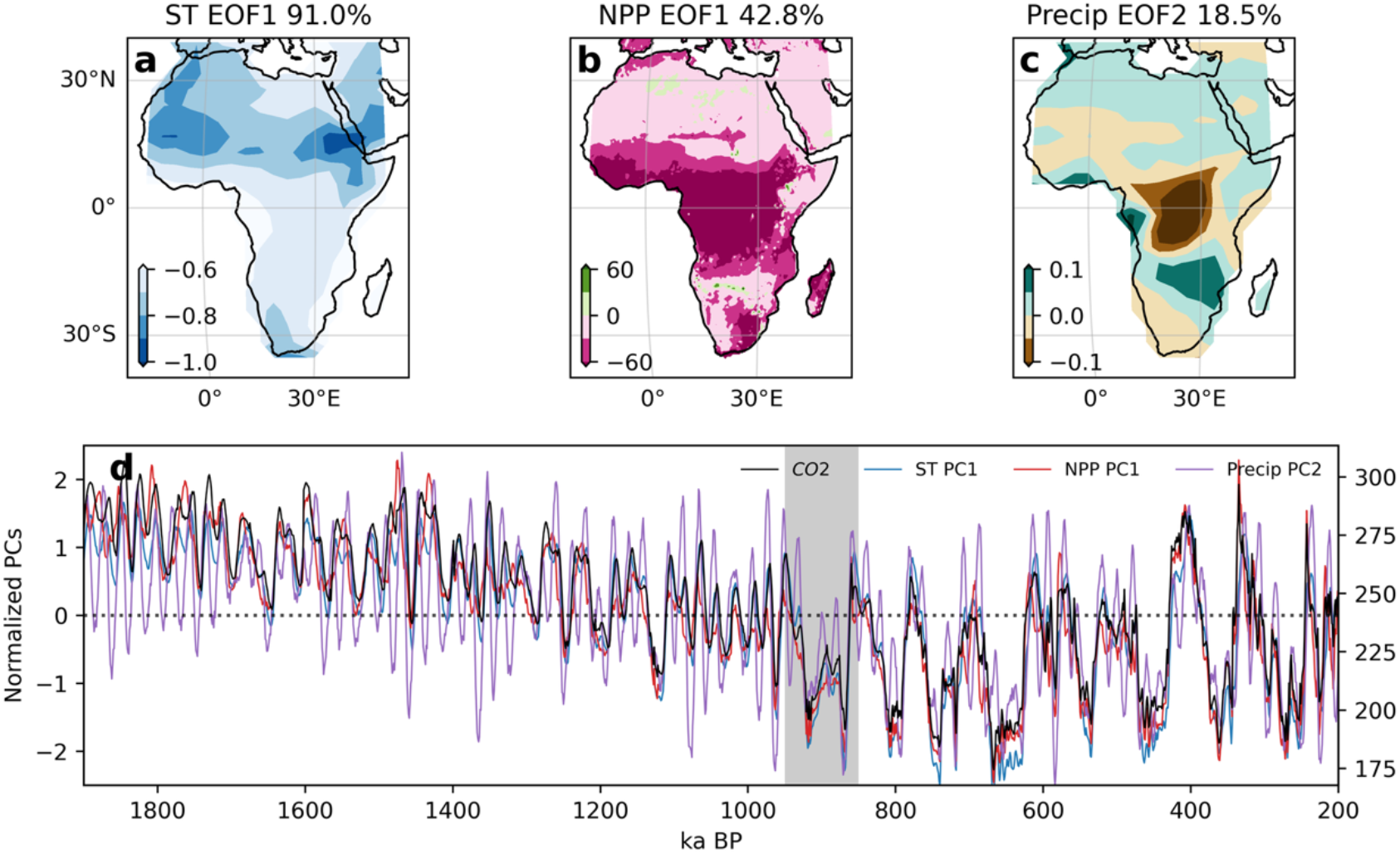
The CO_2_-driven climate components in model simulation. **(a)** Reconstructed surface temperature from the 1^st^ Empirical Orthogonal Function (EOF) analysis of African surface temperature (ST) over the bottleneck event (0.95-0.85 Ma). The percentage is the explained variance of the EOF mode. **(b)** for net primary production. **(c)** for precipitation but with EOF2. **(d)** The normalized principal components of surface temperature (blue), NPP (red), and precipitation (purple), and CO_2_ variations used as one of the forcings in the 3 Ma simulation (brown; right y-axis).

**Figure S6.**
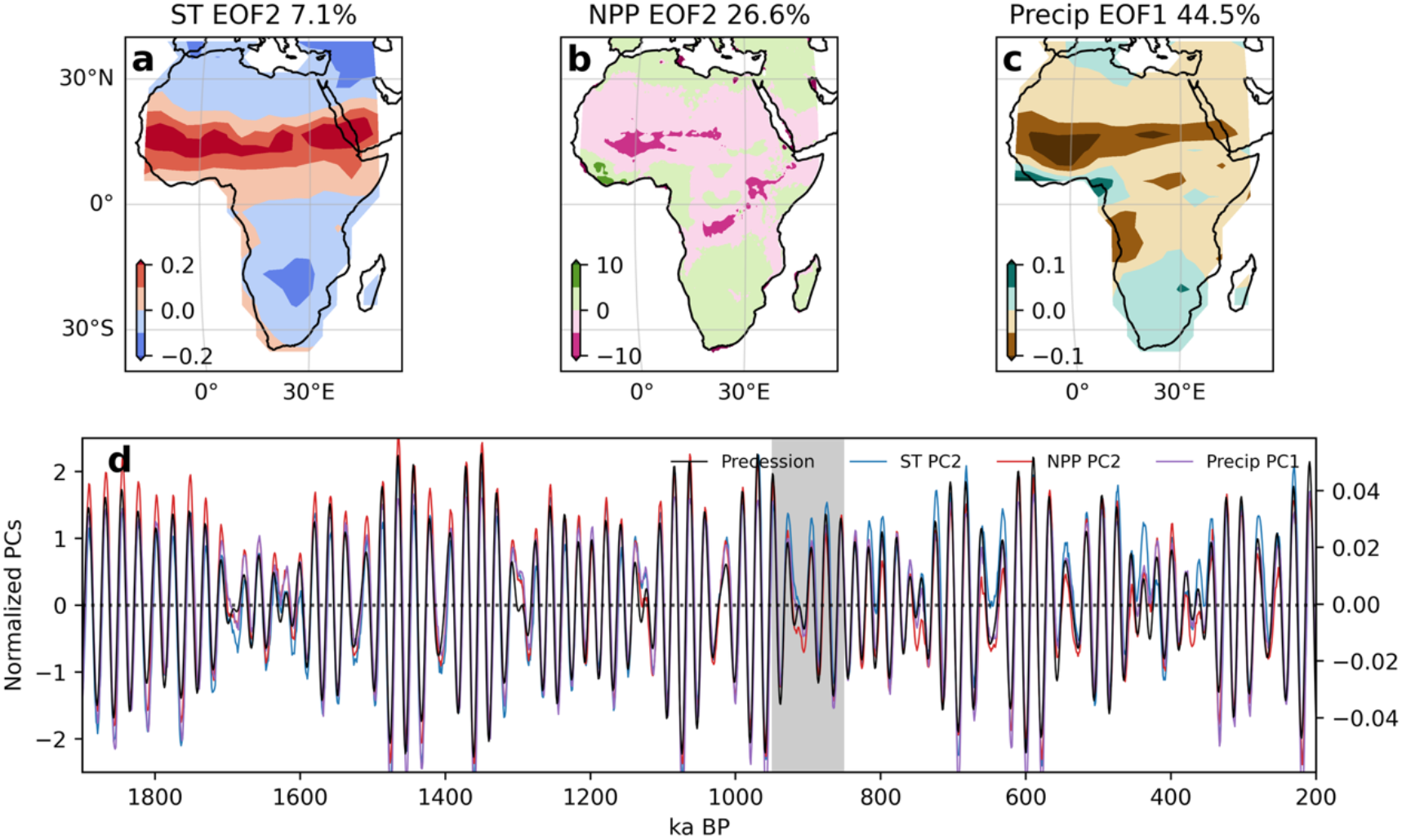
The solar radiation-driven climate components in model simulation. **(a)** Reconstructed surface temperature from the EOF2 of African surface temperature over the bottleneck event (0.95-0.85 Ma). The percentage is the explained variance of the EOF mode. **(b)** for net primary production. **(c)** for precipitation but with EOF1. **(d)** The normalized principal components of surface temperature (blue), NPP (red), and precipitation (purple). Changes in earth’s precession parameter used as one of the forcings in the 3 Ma simulation ^19^ (black; right y-axis).

**Figure S7.**
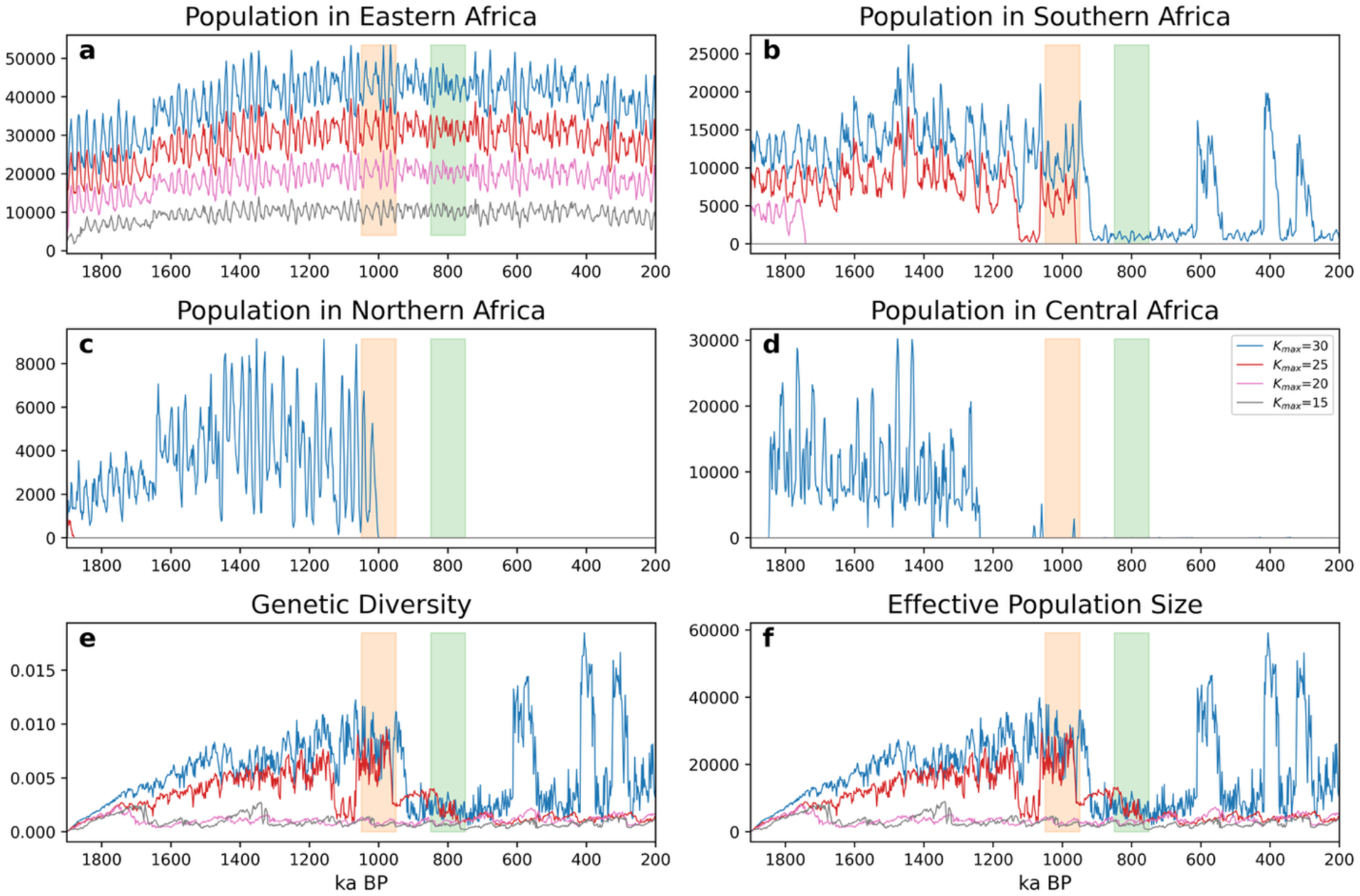
Eastern Africa as a genetic reservoir indicated by its stable survival with lower cultural development. **(a)** Time series of population in eastern Africa for the simulations with K_max_=30 (blue), K_max_=25 (red), K_max_=20 (magenta), and K_max_=15 (gray). **(b)** for southern Africa. **(c)** for northern Africa. **(d)** for central Africa. **(e)** for genetic diversity. **(f)** for effective population size.

**Figure S8.**
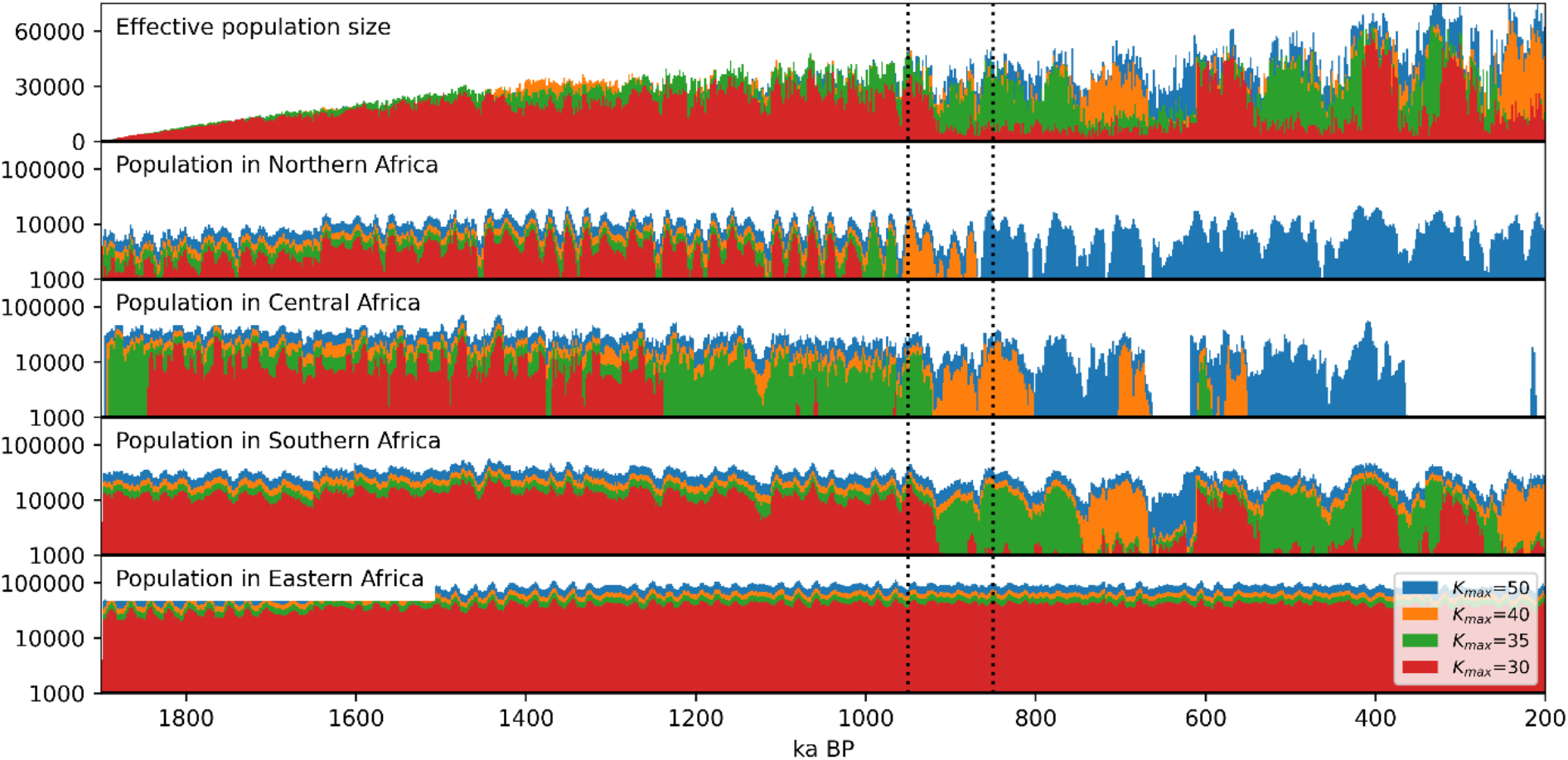
Surviving ability of regions after the MPT. Effective population size for simulations with K_max_=50 (blue), K_max_= 40 (orange), K_max_= 35 (green), K_max_=30 (red) in top. Temporal population in northern Africa (with log scale) and below for central Africa, for southern Africa, for eastern Africa. The dotted lines indicate the bottleneck event (950 ka to 850 ka).

**Figure S9.**
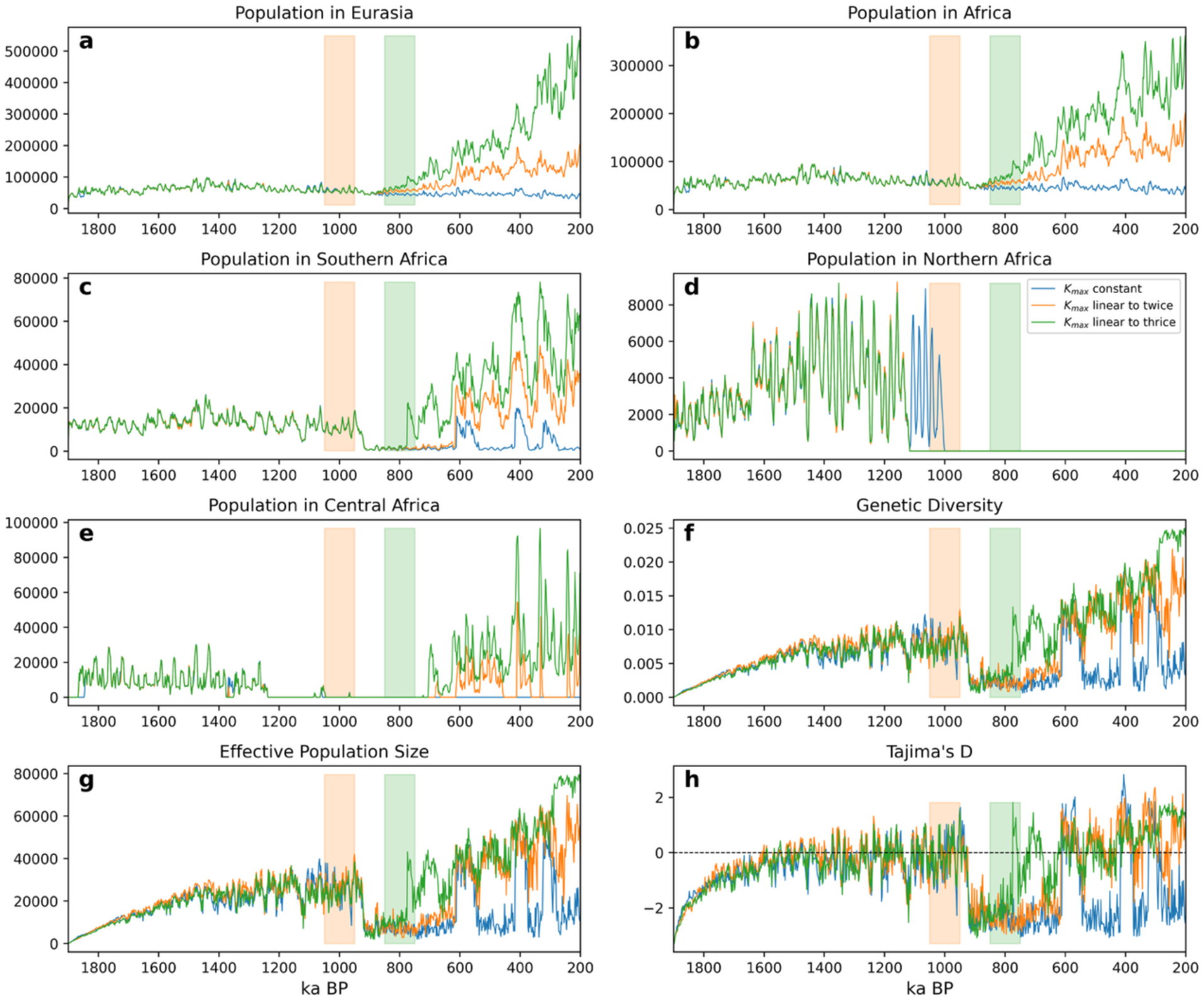
The recovery of DNA diversity after the bottleneck with linear cultural developments. **(a)** Time series of population in Eurasia for the simulations with no K_max_ change (blue), a linear increase of K_max_ to doubled K_max_ after 900 ka (orange), to tripled K_max_ (green). **(b)** for Africa. **(c)** for southern Africa. **(d)** for northern Africa. **(e)** for central Africa. **(f)** for genetic diversity. **(g)** for effective population size. **(h)** for Tajima’s D.

**Figure S10.**
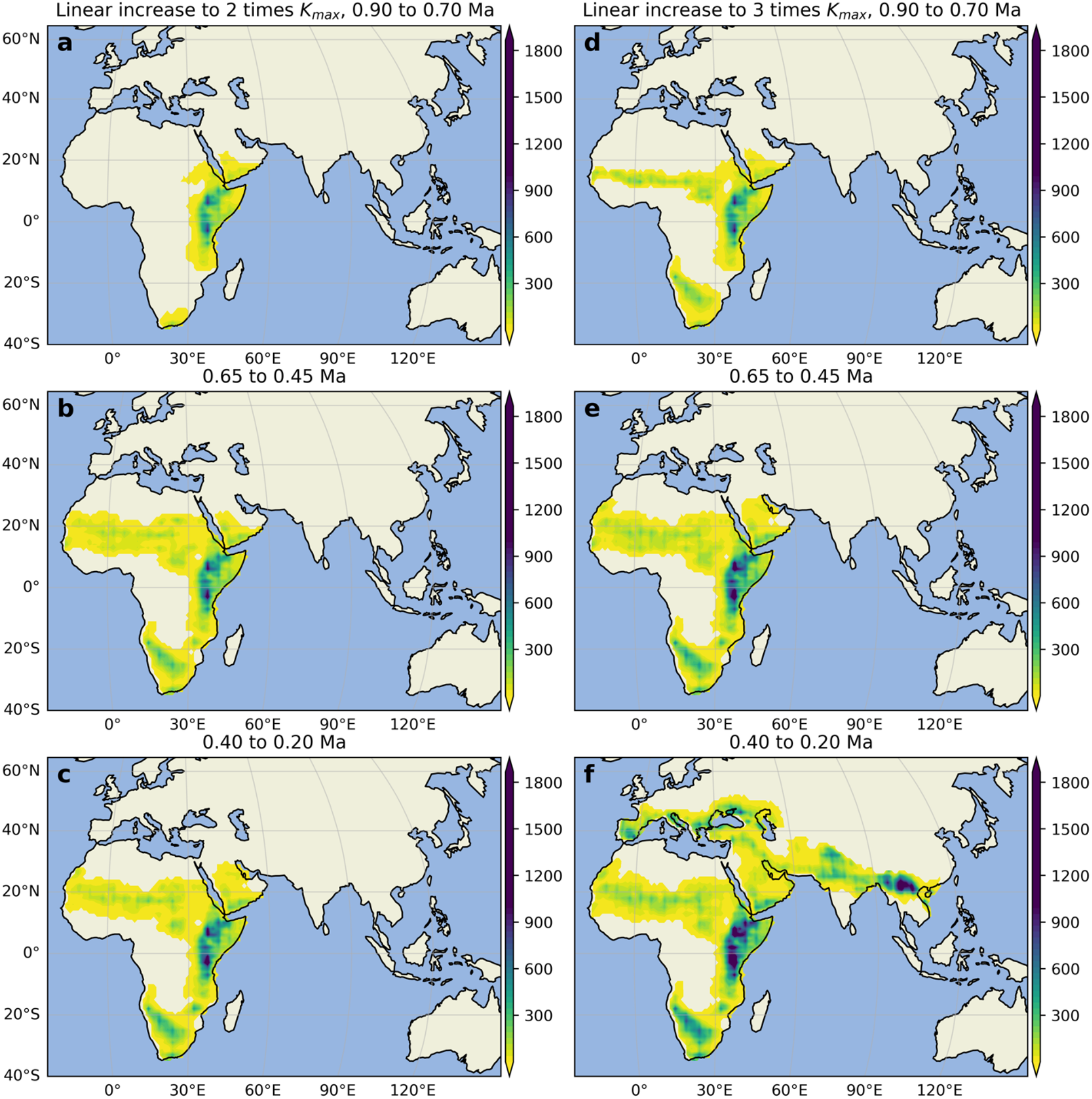
Time slices of population densities of the ABM simulation with linear cultural development to twice and tripled values after the bottleneck. **(a)** Mean population over 0.9 to 0.7 Ma for the simulation with a linear increase of K_max_ to doubled K_max_ after 0.9 Ma. **(b)** for 0.65 to 0.45 Ma. **(c)** 0.4 to 0.2 Ma. **(d)**-**(f)** same as **a** to **c**, but linear increase of K_max_ to tripled K_max_.

---

**Table S1.** Parameters used in the standard agent-based model simulation. The unit of mutation rate is cite per year.

| Variable | Long name | Value |
| --- | --- | --- |
| max_age_f | Maximum age of a female | 55 |
| max_age_m | Maximum age of a male | 55 |
| pm | Movement probability per individual per year | 0.3 |
| fs_m | Fertility starting age of a male | 13 |
| fs_f | Fertility starting age of a female | 11 |
| fe_m | Fertility ending age of a male | 55 |
| fe_f | Fertility ending age of a female | 42 |
| B | Birth rate | 0.12 |
| NPP<sub>w</sub> | A number that determines by how much the availability of water resources increases local NPP | 0.3 |
| NPP<sub>max</sub> | NPP at which carrying capacity reaches its maximum | 900.0 (gC/m<sup>2</sup>) |
| li | Interbirth interval | 2 |
| inx_radius | Interaction radius | 1 |
| lowest_lat | South boundary | -37 |
| lowest_lon | West boundary | -22 |
| largest_lat | North boundary | 62 |
| largest_lon | East boundary | 142 |
| max_age_uncertainty | The range around the max_age parameter in a uniform distribution to randomly select a max_age for each individual | 0.1 |
| $\mu$ | Mutation rate | 2.7e-8 |
| K<sub>max</sub> | Maximum carrying capacity. | 30 |

## Notes

### Competing Interest Statement

The authors have declared no competing interest.

## REFERENCES

1. Timmermann, A., Raia, P., Mondanaro, A., Zollikofer, C.P.E., de Leon, M.P., Zeller, E., and Yun, K.S. (2024). Past climate change effects on human evolution. Nat Rev Earth Env. 10.1038/s43017-024-00584-4.

2. Foerster, V., Asrat, A., Bronk Ramsey, C., Brown, E.T., Chapot, M.S., Deino, A., Duesing, W., Grove, M., Hahn, A., Junginger, A., et al. (2022). Pleistocene climate variability in eastern Africa influenced hominin evolution. Nat Geosci 15, 805–811. 10.1038/s41561-022-01032-y.

3. Zan, J., Louys, J., Dennell, R., Petraglia, M., Ning, W., Fang, X., Zhang, W., and Hu, Z. (2024). Mid-Pleistocene aridity and landscape shifts promoted Palearctic hominin dispersals. Nat Commun 15, 10279. 10.1038/s41467-024-54767-0.

4. Gosling, W.D., Scerri, E.M.L., and Kaboth-Bahr, S. (2022). The climate and vegetation backdrop to hominin evolution in Africa. Philos Trans R Soc Lond B Biol Sci 377, 20200483. 10.1098/rstb.2020.0483.

5. Ragsdale, A.P., Weaver, T.D., Atkinson, E.G., Hoal, E.G., Möller, M., Henn, B.M., and Gravel, S. (2023). A weakly structured stem for human origins in Africa. Nature 617, 755–763. 10.1038/s41586-023-06055-y.

6. Bergström, A., Stringer, C., Hajdinjak, M., Scerri, E.M.L., and Skoglund, P. (2021). Origins of modern human ancestry. Nature 590, 229–237. 10.1038/s41586-021-03244-5.

7. Scerri, E.M.L. (2023). One species, many roots? Nat Ecol Evol 7, 975–976. 10.1038/s41559-023-02080-2.

8. Grundler, M.C., Terhorst, J., and Bradburd, G.S. (2025). A geographic history of human genetic ancestry. Science 387, 1391–1397. 10.1126/science.adp4642.

9. Padilla-Iglesias, C., Xue, Z., Leonardi, M., Paijmans, J.L.A., Colucci, M., Hovhannisyan, A., Maisano-Delser, P., Blanco-Portillo, J., Ioannidis, A.G., Lucarini, G., et al. (2025). Pan-African metapopulation model explains Homo sapiens genetic and morphological evolution. bioRxiv, 2025.2005.2022.655514. 10.1101/2025.05.22.655514.

10. Posth, C., Yu, H., Ghalichi, A., Rougier, H., Crevecoeur, I., Huang, Y., Ringbauer, H., Rohrlach, A.B., Nagele, K., Villalba-Mouco, V., et al. (2023). Palaeogenomics of Upper Palaeolithic to Neolithic European hunter-gatherers. Nature 615, 117–126. 10.1038/s41586-023-05726-0.

11. Salem, N., van de Loosdrecht, M.S., Sumer, A.P., Vai, S., Hubner, A., Peter, B., Bianco, R.A., Lari, M., Modi, A., Al-Faloos, M.F.M., et al. (2025). Ancient DNA from the Green Sahara reveals ancestral North African lineage. Nature 641, 144–150. 10.1038/s41586-025-08793-7.

12. Durvasula, A., and Sankararaman, S. (2020). Recovering signals of ghost archaic introgression in African populations. Sci Adv 6, eaax5097. 10.1126/sciadv.aax5097.

13. Ramachandran, S., Deshpande, O., Roseman, C.C., Rosenberg, N.A., Feldman, M.W., and Cavalli-Sforza, L.L. (2005). Support from the relationship of genetic and geographic distance in human populations for a serial founder effect originating in Africa. Proc Natl Acad Sci U S A 102, 15942–15947. 10.1073/pnas.0507611102.

14. Betti, L., Balloux, F., Amos, W., Hanihara, T., and Manica, A. (2009). Distance from Africa, not climate, explains within-population phenotypic diversity in humans. Proc Biol Sci 276, 809–814. 10.1098/rspb.2008.1563.

15. Raia, P., Mondanaro, A., Melchionna, M., Di Febbraro, M., Diniz, J.A.F., Rangel, T.F., Holden, P.B., Carotenuto, F., Edwards, N.R., Lima-Ribeiro, M.S., et al. (2020). Past Extinctions of Homo Species Coincided with Increased Vulnerability to Climatic Change. One Earth 3, 480–490. 10.1016/j.oneear.2020.09.007.

16. Degroot, D., Anchukaitis, K.J., Tierney, J.E., Riede, F., Manica, A., Moesswilde, E., and Gauthier, N. (2022). The history of climate and society: a review of the influence of climate change on the human past. Environmental Research Letters 17, 103001. 10.1088/1748-9326/ac8faa.

17. deMenocal, P.B., and Stringer, C. (2016). Human migration: Climate and the peopling of the world. Nature 538, 49–50. 10.1038/nature19471.

18. Timmermann, A., Yun, K.S., Raia, P., Ruan, J.Y., Mondanaro, A., Zeller, E., Zollikofer, C., de León, M.P., Lemmon, D., Willeit, M., and Ganopolski, A. (2022). Climate effects on archaic human habitats and species successions. Nature 604, 495–501. 10.1038/s41586-022-04600-9.

19. Jakobsson, M., Bernhardsson, C., McKenna, J., Hollfelder, N., Vicente, M., Edlund, H., Coutinho, A., Sjodin, P., Brink, J., Zipfel, B., et al. (2026). Homo sapiens-specific evolution unveiled by ancient southern African genomes. Nature 650, 156–163. 10.1038/s41586-025-09811-4.

20. Vahdati, A.R., Weissmann, J.D., Timmermann, A., de León, M.P., and Zollikofer, C.P.E. (2022). Exploring Late Pleistocene hominin dispersals, coexistence and extinction with agent-based multi-factor models. Quaternary Sci Rev 279. 10.1016/j.quascirev.2022.107391.

21. Hallett, E.Y., Leonardi, M., Cerasoni, J.N., Will, M., Beyer, R., Krapp, M., Kandel, A.W., Manica, A., and Scerri, E.M.L. (2025). Major expansion in the human niche preceded out of Africa dispersal. Nature 644, 115–121. 10.1038/s41586-025-09154-0.

22. Mondanaro, A., Melchionna, M., Di Febbraro, M., Castiglione, S., Holden, P.B., Edwards, N.R., Carotenuto, F., Maiorano, L., Modafferi, M., Serio, C., et al. (2020). A Major Change in Rate of Climate Niche Envelope Evolution during Hominid History. Iscience 23. 10.1016/j.isci.2020.101693.

23. Wakano, J.Y., Gilpin, W., Kadowaki, S., Feldman, M.W., and Aoki, K. (2018). Ecocultural range-expansion scenarios for the replacement or assimilation of Neanderthals by modern humans. Theoretical Population Biology 119, 3–14. 10.1016/j.tpb.2017.09.004.

24. Zeller, E., Timmermann, A., Yun, K.S., Raia, P., Stein, K., and Ruan, J.Y. (2023). Human adaptation to diverse biomes over the past 3 million years. Science 380, 604–608. 10.1126/science.abq1288.

25. Vahdati, A.R., Weissmann, J.D., Timmermann, A., de León, M.S.P., and Zollikofer, C.P.E. (2019). Drivers of Late Pleistocene human survival and dispersal: an agent-based modeling and machine learning approach. Quaternary Sci Rev 221. 10.1016/j.quascirev.2019.105867.

26. Posth, C., Renaud, G., Mittnik, A., Drucker, D.G., Rougier, H., Cupillard, C., Valentin, F., Thevenet, C., Furtwängler, A., Wissing, C., et al. (2016). Pleistocene Mitochondrial Genomes Suggest a Single Major Dispersal of Non-Africans and a Late Glacial Population Turnover in Europe. Current Biology 26, 827–833. 10.1016/j.cub.2016.01.037.

27. Timmermann, A., Wasay, A., and Raia, P. (2024). Phase synchronization between culture and climate forcing. P Roy Soc B-Biol Sci 291. 10.1098/rspb.2024.0320.

28. Benguigui, M., and Arenas, M. (2014). Spatial and Temporal Simulation of Human Evolution. Methods, Frameworks and Applications. Curr Genomics 15, 245–255. Doi 10.2174/1389202915666140506223639.

29. Yun, K.S., Timmermann, A., Lee, S.S., Willeit, M., Ganopolski, A., and Jadhav, J. (2023). A transient coupled general circulation model (CGCM) simulation of the past 3 million years. Clim Past 19, 1951–1974. 10.5194/cp-19-1951-2023.

30. Lankheet, I., Chowdhury, A., Tellgren-Roth, C., Jolly, C., Soares, A.E.R., de Navascues, M., Pacchiarotti, S., Maselli, L., Kouarata, G., Donzo, J.P., et al. (2026). Revisiting the African mtDNA landscape through complete mitochondrial genomes. Commun Biol 9. 10.1038/s42003-026-10330-9.

31. Tommasi, A., Boscolo Agostini, R., Villani, G., Rambaldi Migliore, N., Vizzari, M.T., Cardinali, I., Di Gerlando, R., Nicolini, V., Sorasio, G., Santos, P., et al. (2025). Fifteen millennia of human mitogenome evolution in Sicily. Sci Adv 11, eady1674. 10.1126/sciadv.ady1674.

32. Charlesworth, B. (2009). Effective population size and patterns of molecular evolution and variation. Nat Rev Genet 10, 195–205. 10.1038/nrg2526.

33. Willeit, M., Ganopolski, A., Calov, R., and Brovkin, V. (2019). Mid-Pleistocene transition in glacial cycles explained by declining CO2 and regolith removal. Sci Adv 5. 10.1126/sciadv.aav7337.

34. Lüthi, D., Le Floch, M., Bereiter, B., Blunier, T., Barnola, J.M., Siegenthaler, U., Raynaud, D., Jouzel, J., Fischer, H., Kawamura, K., and Stocker, T.F. (2008). High-resolution carbon dioxide concentration record 650,000-800,000 years before present. Nature 453, 379–382. 10.1038/nature06949.

35. Berger, A., and Loutre, M.F. (1991). Insolation Values for the Climate of the Last 10000000 Years. Quaternary Sci Rev 10, 297–317. Doi 10.1016/0277-3791(91)90033-Q.

36. Center, N.G.D. (1993). 5-minute Gridded Global Relief Data (ETOPO5).

37. Tamura, K., and Nei, M. (1993). Estimation of the Number of Nucleotide Substitutions in the Control Region of Mitochondrial-DNA in Humans and Chimpanzees. Mol Biol Evol 10, 512–526. 10.1093/oxfordjournals.molbev.a040023.

38. Bouckaert, R., Vaughan, T.G., Barido-Sottani, J., Duchêne, S., Fourment, M., Gavryushkina, A., Heled, J., Jones, G., Kühnert, D., De Maio, N., et al. (2019). BEAST 2.5: An advanced software platform for Bayesian evolutionary analysis. Plos Computational Biology 15. 10.1371/journal.pcbi.1006650.

39. Watterson, G.A. (1975). On the number of segregating sites in genetical models without recombination. Theor Popul Biol 7, 256–276. 10.1016/0040-5809(75)90020-9.

40. Suchard, M.A., Lemey, P., Baele, G., Ayres, D.L., Drummond, A.J., and Rambaut, A. (2018). Bayesian phylogenetic and phylodynamic data integration using BEAST 1.10. Virus Evol 4. 10.1093/ve/vey016.

41. Tavaré, S. (1986). Some probabilistic and statistical problems on the analysis of DNA sequences. Lectures on Mathematics in the Life Sciences 17, 57–86.

42. Drummond, A.J., Rambaut, A., Shapiro, B., and Pybus, O.G. (2005). Bayesian coalescent inference of past population dynamics from molecular sequences. Mol Biol Evol 22, 1185–1192. 10.1093/molbev/msi103.

43. Rambaut, A., Drummond, A.J., Xie, D., Baele, G., and Suchard, M.A. (2018). Posterior Summarization in Bayesian Phylogenetics Using Tracer 1.7. Syst Biol 67, 901–904. 10.1093/sysbio/syy032.

44. Head, M.J., and Gibbard, P.L. (2005). Early-Middle Pleistocene transitions: An overview and recommendation for the defining boundary. Geol Soc Spec Publ 247, 1–18. Doi 10.1144/Gsl.Sp.2005.247.01.01.

45. Clark, P.U., and Pollard, D. (1998). Origin of the middle Pleistocene transition by ice sheet erosion of regolith. Paleoceanography 13, 1–9. Doi 10.1029/97pa02660.

46. deMenocal, P.B., and Bloemendal, J. (1995). Plio-Pleistocene climatic variability in subtropical Africa and the paleoenvironment of hominid evolution: A combined data-model approach. Paleoclimate and Evolution, with Emphasis on Human Origins, 262–288.

47. Tiedemann, R., Sarnthein, M., and Shackleton, N.J. (1994). Astronomic Timescale for the Pliocene Atlantic Delta-O-18 and Dust Flux Records of Ocean Drilling Program Site-659. Paleoceanography 9, 619–638. Doi 10.1029/94pa00208.

48. Grant, K.M., Rohling, E.J., Westerhold, T., Zabel, M., Heslop, D., Konijnendijk, T., and Lourens, L. (2017). A 3 million year index for North African humidity/aridity and the implication of potential pan-African Humid periods. Quaternary Sci Rev 171, 100–118. 10.1016/j.quascirev.2017.07.005.

49. Lisiecki, L.E., and Raymo, M.E. (2005). A Pliocene-Pleistocene stack of 57 globally distributed benthic δ18O records. Paleoceanography 20. 10.1029/2005pa001164.

50. Gallotti, R., and Mussi, M. (2018). The Emergence of the Acheulean in East Africa: Historical Perspectives and Current Issues. Vertebr Paleobiol Pa, 1–12. 10.1007/978-3-319-75985-2_1.

51. Paige, J., and Perreault, C. (2024). 3.3 million years of stone tool complexity suggests that cumulative culture began during the Middle Pleistocene. P Natl Acad Sci USA 121. 10.1073/pnas.2319175121.

52. Deng, Y., Nielsen, R., and Song, Y.S. (2024). A previously reported bottleneck in human ancestry 900 kya is likely a statistical artifact. bioRxiv, 2024.2010.2001.615851. 10.1101/2024.10.01.615851.

53. Hu, W.J., Hao, Z.Q., Du, P.Y., Di Vincenzo, F., Manzi, G., Cui, J.L., Fu, Y.X., Pan, Y.H., and Li, H.P. (2023). Genomic inference of a severe human bottleneck during the Early to Middle Pleistocene transition. Science 381, 979–984. 10.1126/science.abq7487.

54. Diniz, J.A.F., Soares, T.N., Lima, J.S., Dobrovolski, R., Landeiro, V.L., Telles, M.P.D., Rangel, T.F., and Bini, L.M. (2013). Mantel test in population genetics. Genet Mol Biol 36, 475–485. Doi 10.1590/S1415-47572013000400002.

55. Hannachi, A., Jolliffe, I.T., and Stephenson, D.B. (2007). Empirical orthogonal functions and related techniques in atmospheric science: A review. Int J Climatol 27, 1119–1152. 10.1002/joc.1499.

56. Jakobsson, M., Bernhardsson, C., McKenna, J., Hollfelder, N., Vicente, M., Edlund, H., Coutinho, A., Sjodin, P., Brink, J., Zipfel, B., et al. (2025). Homo sapiens-specific evolution unveiled by ancient southern African genomes. Nature. 10.1038/s41586-025-09811-4.

57. Kolodny, O., Creanza, N., and Feldman, M.W. (2016). Game-Changing Innovations: How Culture Can Change the Parameters of Its Own Evolution and Induce Abrupt Cultural Shifts. Plos Comput Biol 12. 10.1371/journal.pcbi.1005302.

58. Shao, Y.P., Wegener, C., Klein, K., Schmidt, I., and Weniger, G.C. (2024). Reconstruction of human dispersal during Aurignacian on pan-European scale. Nat Commun 15. 10.1038/s41467-024-51349-y.

59. Harvati, K., Röding, C., Bosman, A.M., Karakostis, F.A., Grün, R., Stringer, C., Karkanas, P., Thompson, N.C., Koutoulidis, V., Moulopoulos, L.A., et al. (2019). Apidima Cave fossils provide earliest evidence of Homo sapiens in Eurasia. Nature 571, 500–504. 10.1038/s41586-019-1376-z.

60. Hershkovitz, I., Weber, G.W., Quam, R., Duval, M., Grün, R., Kinsley, L., Ayalon, A., Bar-Matthews, M., Valladas, H., Mercier, N., et al. (2018). The earliest modern humans outside Africa. Science 359, 456–459. 10.1126/science.aap8369.

61. Rito, T., Vieira, D., Silva, M., Conde-Sousa, E., Pereira, L., Mellars, P., Richards, M.B., and Soares, P. (2019). A dispersal of Homo sapiens from southern to eastern Africa immediately preceded the out-of-Africa migration. Sci Rep-Uk 9. 10.1038/s41598-019-41176-3.

62. Mounier, A., and Lahr, M.M. (2019). Deciphering African late middle Pleistocene hominin diversity and the origin of our species. Nat Commun 10. 10.1038/s41467-019-11213-w.

63. Scerri, E.M.L., Thomas, M.G., Manica, A., Gunz, P., Stock, J.T., Stringer, C., Grove, M., Groucutt, H.S., Timmermann, A., Rightmire, G.P., et al. (2018). Did Our Species Evolve in Subdivided Populations across Africa, and Why Does It Matter? Trends in Ecology & Evolution 33, 582–594. 10.1016/j.tree.2018.05.005.

64. Zhou, S., Zhou, Y., Di Vincenzo, F., Manzi, G., Hu, W., Hao, Z., Pan, Y.-H., and Li, H. (2025). Ancient human extinction risk: Implications from high-precision computation and fossil evidence. bioRxiv, 2025.2002.2001.636025. 10.1101/2025.02.01.636025.

65. Terhorst, J. (2024). Accelerated Bayesian inference of population size history from recombining sequence data. bioRxiv, 2024.2003.2025.586640. 10.1101/2024.03.25.586640.

66. Cousins, T., and Durvasula, A. (2024). Insufficient evidence for a severe bottleneck in humans during the Early to Middle Pleistocene transition. bioRxiv, 2024.2010.2021.619456. 10.1101/2024.10.21.619456.

67. Gowlett, J.A.J., and Wrangham, R.W. (2013). Earliest fire in Africa: towards the convergence of archaeological evidence and the cooking hypothesis. Azania 48, 5–30. 10.1080/0067270x.2012.756754.

68. Du, A., Zipkin, A.M., Hatala, K.G., Renner, E., Baker, J.L., Bianchi, S., Bernal, K.H., and Wood, B.A. (2018). Pattern and process in hominin brain size evolution are scale-dependent. Proceedings of the Royal Society B-Biological Sciences 285. 10.1098/rspb.2017.2738.

69. Leigh, S.R. (2012). Brain Size Growth and Life History in Human Evolution. Evol Biol 39, 587–599. 10.1007/s11692-012-9168-5.

70. Herbert, T.D., Peterson, L.C., Lawrence, K.T., and Liu, Z.H. (2010). Tropical Ocean Temperatures Over the Past 3.5 Million Years. Science 328, 1530–1534. 10.1126/science.1185435.

71. Margari, V., Hodell, D.A., Parfitt, S.A., Ashton, N.M., Grimalt, J.O., Kim, H., Yun, K.S., Gibbard, P.L., Stringer, C.B., Timmermann, A., and Tzedakis, P.C. (2023). Extreme glacial cooling likely led to hominin depopulation of Europe in the Early Pleistocene. Science 381, 693–698. 10.1126/science.adf4445.

72. Ambrose, S.H. (1998). Late Pleistocene human population bottlenecks, volcanic winter, and differentiation of modern humans. J Hum Evol 34, 623–651. DOI 10.1006/jhev.1998.0219.

73. Schlebusch, C.M., Sjödin, P., Breton, G., Günther, T., Naidoo, T., Hollfelder, N., Sjöstrand, A.E., Xu, J.Z., Gattepaille, L.M., Vicente, M., et al. (2020). Khoe-San Genomes Reveal Unique Variation and Confirm the Deepest Population Divergence in. Mol Biol Evol 37, 2944–2954. 10.1093/molbev/msaa140.

74. Fang, S.-W. (2025). Data and Codes of “A Climate-Driven Collapse in African Human Genomic Diversity ∼900 kyrs Ago”. 10.5281/zenodo.15233604.

